# Predicting the macrovascular contribution to resting-state fMRI functional connectivity at 3 Tesla: A model-informed approach

**DOI:** 10.1101/2024.02.13.580143

**Authors:** Xiaole Z. Zhong, Jonathan R. Polimeni, J. Jean Chen

**Affiliations:** Rotman Research Institute at Baycrest, Toronto, ON, Canada; Department of Medical Biophysics, University of Toronto, Toronto, ON, Canada; Athinoula A. Martinos centre for Biomedical Imaging, Massachusetts General Hospital, Charlestown, MA, USA; Department of Radiology, Harvard Medical School, Boston, MA, USA; Harvard-MIT Program in Health Sciences and Technology, Massachusetts Institute of Technology, Cambridge, MA, USA; Department of Biomedical Engineering, University of Toronto, Toronto, ON, Canada

## Abstract

Macrovascular biases have been a long-standing challenge for fMRI, limiting its ability to detect spatially specific neural activity. Recent experimental studies, including our own (Huck et al., 2023; Zhong et al., 2023), found substantial resting-state macrovascular BOLD fMRI contributions from large veins and arteries, extending into the perivascular tissue at 3 T and 7 T. The objective of this study is to demonstrate the feasibility of predicting, using a biophysical model, the experimental resting-state BOLD fluctuation amplitude (RSFA) and associated functional connectivity (FC) values at 3 Tesla. We investigated the feasibility of both 2D and 3D infinite-cylinder models as well as macrovascular anatomical networks (mVANs) derived from angiograms. Our results demonstrate that: 1) with the availability of mVANs, it is feasible to model macrovascular BOLD FC using both the mVAN-based model and 3D infinite-cylinder models, though the former performed better; 2) biophysical modelling can accurately predict the BOLD pairwise correlation near to large veins (with R^2^ ranging from 0.53 to 0.93 across different subjects), but not near to large arteries; 3) compared with FC, biophysical modelling provided less accurate predictions for RSFA; 4) modelling of perivascular BOLD connectivity was feasible at close distances from veins (with R^2^ ranging from 0.08 to 0.57), but not arteries, with performance deteriorating with increasing distance. While our current study demonstrates the feasibility of simulating macrovascular BOLD in the resting state, our methodology may also apply to understanding task-based BOLD. Furthermore, these results suggest the possibility of correcting for macrovascular bias in resting-state fMRI and other types of fMRI using biophysical modelling based on vascular anatomy.

## 1 Introduction

The use of resting-state blood-oxygenation level-dependent (BOLD) functional magnetic resonance imaging (rs-fMRI) has become increasingly valuable in assessing brain health and mapping brain connectivity (Srivastava et al., 2022; van den Heuvel & Hulshoff Pol, 2010). There has been extensive use of this technique in the study of numerous neurological conditions (Ovadia-Caro et al., 2014; Vemuri et al., 2012), as well as in the study of brain development and ageing processes (Andrews-Hanna et al., 2007; Geerligs et al., 2015 (Andrews-Hanna et al., 2007; Geerligs et al., 2015). The most common method for calculating functional-connectivity (FC) using resting-state magnetic resonance imaging (rs-fcMRI) is Pearson’s correlation analysis (Bandettini et al., 1993; Biswal et al., 1995), in which the BOLD signals from seed voxels are correlated with those from the rest of the brain. Commonly, correlation is interpreted as reflecting synchronous neuronal activity (Biswal et al., 1995; Pan et al., 2015). However, the contribution of macrovascular networks to functional connectivity (FC) in grey matter (GM) has been demonstrated in our previous study (Zhong et al., 2023), which indicated that, at 3 Tesla, such effects occurred not only within the confines of macrovasculature but also more than 10 mm away from individual microvascular blood vessels. Similar results were also shown in a recent publication by Huck et al. (Bernier et al., 2021; Huck et al., 2023), which suggested that at 7 T, large veins could significantly bias the rs-fMRI metrics (including low-frequency fluctuation amplitude, regional connectivity inhomogeneity, Hurst exponent, and eigenvector centrality). However, there is still very limited systematic investigation into macrovascular contributions to rs-fMRI.

Unlike in rs-fMRI, the question of vein versus brain has been debated for many years in task-based fMRI. Boxerman and colleagues demonstrated, through both rodent experiments and simulation studies, that macrovasculature affects gradient-recalled-echo (GRE) BOLD contrast more than microvasculature (Boxerman et al., 1995). Despite the fact that microvascular sensitivity increases with an increase in the main magnetic field (Menon & Goodyear, 1999), veins still account for the majority of the contribution to GRE BOLD signals at 7T (Menon, 2012; UIudag et al., 2009). Previous studies in rats showed that venous structure could be detected using rs-fcMRI (Hyde and Li, 2014), and that the BOLD response from veins is nearly double that from microvessels in brain tissue at 11.7 T (Yu et al., 2012). Moreover, several studies have suggested that even penetrating arteries and ascending veins are also not tightly coupled with local neural activity (Uludağ & Blinder, 2018), despite their much smaller size relative to the macrovasculature investigated in this study.

There have also been numerous approaches developed to remove the venous BOLD effect. The simplest method of correcting the bias appears to be to mask out the detectable macrovasculature using vascular masks, such as segmented from angiograms. However, as work by ourselves and others have shown (Huck et al., 2023; Zhong et al., 2023)(Bernier et al., 2021; Huck et al., 2023)(Huck et al., 2023; Zhong et al., 2023), macrovascular effects can extend well into the perivascular tissue, making masking-based correction inadequate. Spin-echo (SE) BOLD has also been proposed as an alternative to the conventional GRE BOLD to suppress the strong macrovascular susceptibility effects. Nevertheless, this option sacrifices signal-to-noise ratio (Menon, 2012), and the use of typical echo-planar imaging (EPI) readouts even with SE refocusing is still susceptible to large-vein T2* effects (Goense & Logothetis, 2006; Ragot & Chen, 2019). The BOLD image phase has also been proposed as a regressor to remove macrovascular effects in task-based fMRI (Menon, 2002; Stanley et al., 2021), assuming that the signal magnitude and phase scale with one another, and that the extravascular (EV) contribution is dwarfed by the intravascular (IV) contribution due to low voxel resolution at conventional field strengths, which may not always be the case. As vascular diameter increases, it becomes less appropriate to ignore the IV phase offsets and vascular orientation effects, as discussed in our previous work (Zhong & Chen, 2022, 2023). Most recently, Huck et al. (Huck et al., 2023) advocated the use of high-order polynomials for modelling the fall-off of extravascular fields over space and subsequently removing venous biases, both proximal and distal to the vasculature. However, the authors concluded that their model is not sufficient to correct venous bias in rs-fMRI data. In light of these recent findings, it may be necessary to consider a more comprehensive modelling approach for the vascular BOLD contributions as the initial step in correcting for them.

The modelling of macrovascular contributions to grey matter (GM) FC with a closed-form analytical model is a challenging undertaking. Such modelling ideally utilises parameters such as vascular diameter, vascular orientation, and blood oxygenation, some of which are difficult to measure reliably in the human brain. Early work by Ogawa et al. on BOLD biophysical models demonstrated that the BOLD signal is dependent on the orientation and diameter of the vessels, as well as blood oxygenation (Ogawa, Menon, et al., 1993). The dependences of BOLD signals on the vascular geometry (mainly the large pial veins) were also evident in the biophysical simulations based on realistic vascular anatomy (Gagnon et al., 2015) and in in-vivo fMRI measurements (Fracasso et al., 2018; Viessmann et al., 2019) as well as in our previous simulations (Zhong & Chen, 2022). Further simulations also revealed that the position of a large vessel position within an fMRI voxel plays a significant role in modulating macrovascular BOLD (Sedlacik et al., 2007), increasing the complexity of modelling macrovascular effects (Zhong & Chen, 2023). Despite not having been investigated previously, other factors (such as patient positioning and spatial resolution) may also have a significant impact on modelling macrovascular effects. Instead of using a closed-form analytical model, it is possible to use a numerical model for calculating macrovascular BOLD signals that may simplify these issues. A kernel-based approach, proposed in previous studies (Cheng et al., 2009; Salomir et al., 2003), involves convolving kernels (representing point-spread functions) with susceptibility maps to compute local offsets in the magnetic field and, consequently, to generate a BOLD signal based on the blood susceptibility. The approach allows us to directly simulate the BOLD signal arising from macrovascular anatomical networks (mVANs), although with the limitation that blood oxygenation remains unknown. It is important to note, however, that this numerical approach will result in approximation errors and require a significant amount of computational resources (as discussed in more detail in the Discussion section). In this regard, it remains to be seen whether the numerical approach has an advantage over the analytical approach.

Though previous attempts at modelling and removal of macrovascular contributions were somewhat effective, they did not fully reveal the various aspects of how the macrovascular anatomy impact the BOLD signal. Motivated by this, the current study is intended to provide a detailed analysis of the macrovascular contribution to FC using a biophysical model. Moreover, two analytical models will be provided in addition to a numerical model in order to provide theoretical understanding and investigate the possibility of simplifying the model. Furthermore, the investigation would be extended to the perivascular tissue in order to gain a better understanding of the probability of modelling the macrovascular contribution beyond the confines of the macrovasculature. As of the time of writing, this is the first study to investigate the modelling of macrovascular origin of FC estimates using biophysical models.

## 2 Method

### 2.1 Data set

In this work, we will compare experimental rs-fMRI data with simulated rs-fMRI data on a voxel-wise basis. We used data selected from the Midnight Scan Club (MSC) dataset (Gordon et al., 2017). The study protocol was approved by the Human Studies Committee and Institute Review Board at Washington University School of Medicine in accordance with the Declaration of Helsinki. This data can be obtained from the OpenNeuro database, with accession number ds000224. Due to the high computational demand of our modelling approach, we balanced computational cost with the generalizability of the results, selecting data from four healthy right-handed young participants, two males and two females, ages 28-34.

### 2.2 MRI acquisition

Each subject underwent 12 imaging sessions on a Siemens TRIO 3T MRI scanner (Siemens Healthcare GmbH, Erlangen, Germany) on separate days. Here, only the protocols related to this study are listed.

In total, 12 2D time-of-flight (TOF) angiograms were acquired, including 4 ascending (transverse, 0.6 × 0.6 × 1.0 mm^3^, 44 slices, TR = 25 ms, TE = 3.34 ms, flip angle (α) = 20°), four left-right encoded, (sagittal, 0.8 × 0.8 × 2.0 mm^3^ thickness, 120 slices, TR = 27 ms, TE = 7.05 ms, α = 60°) and four anterior-posterior encoded (coronal, 0.7 × 0.7 × 2.5 mm^3^ thickness, 128 slices, TR = 28 ms, TE = 7.18 ms, α = 60 degrees) data sets were included. Resting-state fMRI data were acquired with a gradient-recalled-echo BOLD-weighted EPI sequence (TR = 2.2 s, TE = 27 ms, α = 90°, 4 mm isotropic resolution, 36 slices, scan time = 30 minutes). T1-weighted anatomical MPRAGE data were included as well (sagittal, 224 slices, 0.8 mm isotropic resolution, TE = 3.74 ms, TR = 2400 ms, TI = 1000 ms, α = 8 degrees). The participants were instructed to fixate on a white crosshair on a black background during the rs-fMRI scans. An EyeLink 1000 eye-tracking system (SR-Research, Ottawa, Canada, http://www.sr-research.com) was used to monitor participants to determine whether they had any prolonged eye closures, which may indicate sleepiness. There was only one participant who showed prolonged eye closures during the scans.

### 2.3 fMRI processing and analysis

A summary of the rs-fMRI processing procedures can be found in **Figure 1a**. fMRI preprocessing pipeline was implemented with tools from FSL (Jenkinson et al., 2012), AFNI (Cox, 1996) and FreeSurfer (Fischl, 2012). The following steps were included in the preprocessing steps: (a) 3D motion correction (FSL MCFLIRT), (b) slice-timing correction (FSL slicetimer), (c) brain extraction (FSL bet2 and FreeSurfer mri_watershed), (d) rigid body coregistration of functional data to the individual T1 image (FSL FLIRT), (e) regression of the mean signals from white-matter (WM) and cerebrospinal fluid (CSF) regions (fsl_glm), (f) bandpass filtering to obtain frequency band 0.01-0.1 Hz (AFNI 3dBandpass), and (g) spatial smoothing with a 6 mm full-width half-maximum (FWHM) Gaussian kernel (FSL fslmaths). For each participant, after preprocessing, we calculated the voxelwise venous-venous, arterial-arterial, and arterial-venous FC metrics based on the maximum cross-correlation coefficients for the voxels with positive correlation coefficients and the minimum cross-correlation coefficients for the voxels with negative correlation coefficients. In spite of the fact that most fMRI connectivity analysis is based on Pearson’s correlations, we used cross-correlations in order to compare the theoretically predicted correlation coefficients directly with the experimentally measured ones without taking into account time lags.

**Figure 1.**
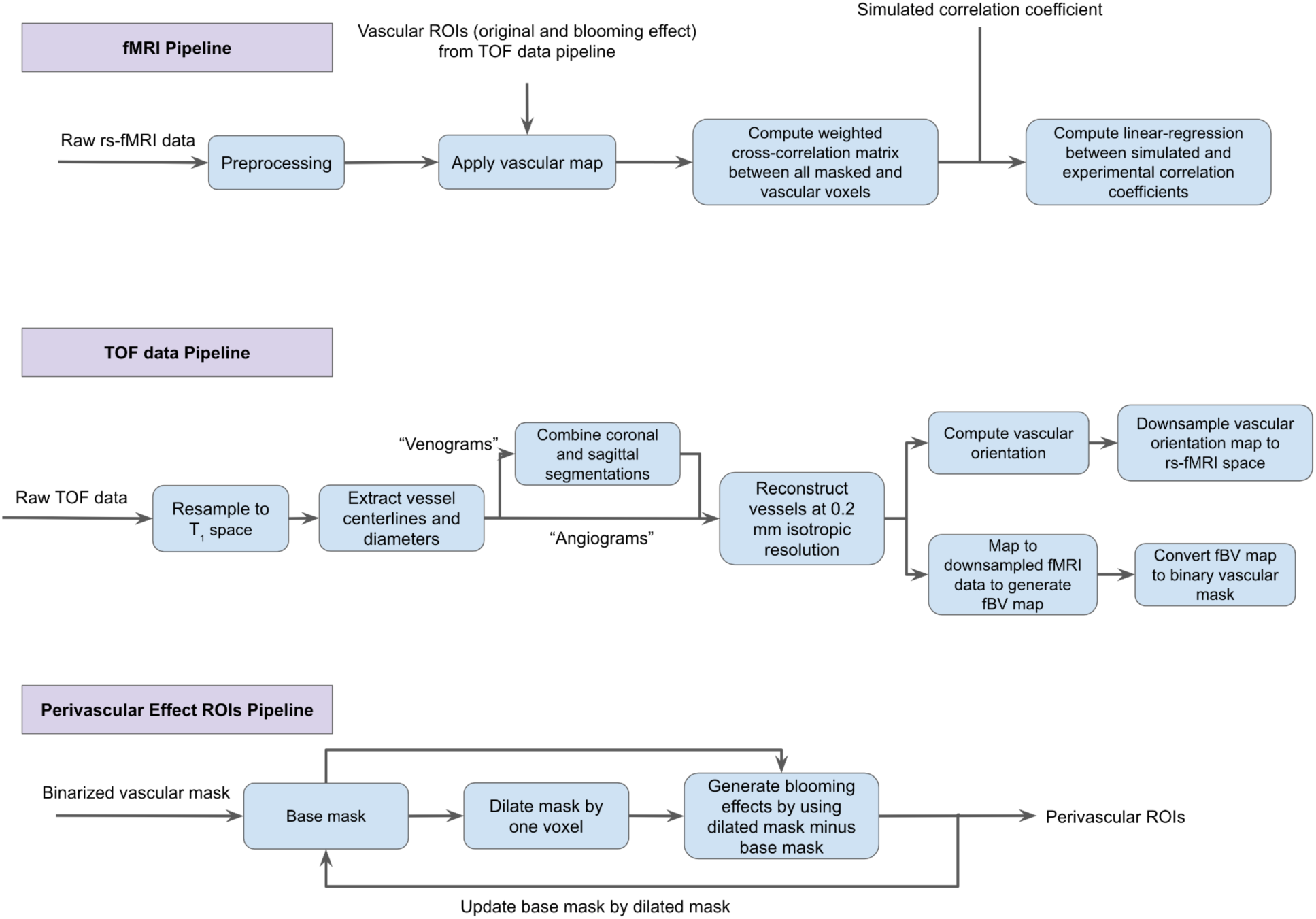
Overview of in-vivo data analysis procedure. a) the fMRI data analysis pipeline; b) the TOF data preprocessing and segmentation pipeline; c) the analysis pipeline for assessing the extravascular effects on rs-fMRI metrics.

### 2.4 Macrovascular segmentation and data processing

The strategies for macrovasculature segmentation and processing are summarised in **Figure 1b**. TOF images were registered to T1 space (FSL MCFLIRT) and segmented using the Braincharter Toolbox (https://github.com/braincharter/vasculature) (Bernier et al., 2018). Visual inspection was performed to ensure the absence of major artefacts. The centreline image encoded with diameter information (“centredia”output from the Braincharter Toolbox) obtained from the raw images (at 0.8 mm isotropic voxel resolution) were pooled from all TOF images across all encoding directions, and where vessel overlaps were detected, the highest diameter estimate is assumed for the overlapping vessel. This combinatorial approach maximised the completeness and signal-to-noise ratio of the resulting mVANs. The combined centrelines were then upsampled to a resolution of 0.2 mm using AFNI (Cox, 1996) with nearest-neighbour interpolation (3dresample). Based on each vascular centreline, a line was constructed that connects two vascular voxels that both neighbour the central voxel in a 9×9×9 voxel matrix. Vascular orientation was calculated as the angle between this line and the z-axis (zenith) and x-axis (azimuth). The zenith angle is further corrected by applying position metrics in MRI header files to compensate for the difference between the participants’ slice direction and the B_0_ direction (scanner z-axis).

Vascular blood-volume fraction (fBV) and orientation maps were manually registered to BOLD space using coordinates from AFNI’s volume selection function (3dAutobox). fBV was estimated by counting the number of high-resolution voxels occupied by vessels within each downsampled voxel, and macrovascular maps were derived from binarized fBV maps. In order to ensure the accuracy of registration, necessary quality control measures have been added. Due to the fact that all TOF images contained both arteries and veins, each arterial and venous map was manually separated after downsampling. Lastly, the measured voxelwise values of orientation and fBV are used to create different simulated voxels containing blood vessels, described in the following. For in-vivo analysis, arterial masks and venous masks were generated by combining fBV maps in rs-fMRI space.

### 2.5 Simulations

All simulation parameters and the values used in all models (**Fig. 2**) are listed in Table 1. In the remainder of this report, the tabulated value for each parameter is assumed unless otherwise stated. It was not expected that results would be affected by the choice of specific *Y* for arteries, veins, and tissues. As the spatial extents of B_0_ inhomogeneities generated by large vessels are much greater than the diffusivity of a water molecule at body temperature (Jensen et al., 2005), we assumed negligible diffusion effects. The simulated BOLD scan parameter values (for example, TR and TE) were adjusted to match the parameter values used in the rs-fMRI scans. At these settings, the arterial inflow effect is deemed negligible (Gao et al., 1996). The Y-dependent T_2_ of tissue and blood were calculated according to previous relaxometry studies (UIudag et al., 2009).

**Figure 2.**
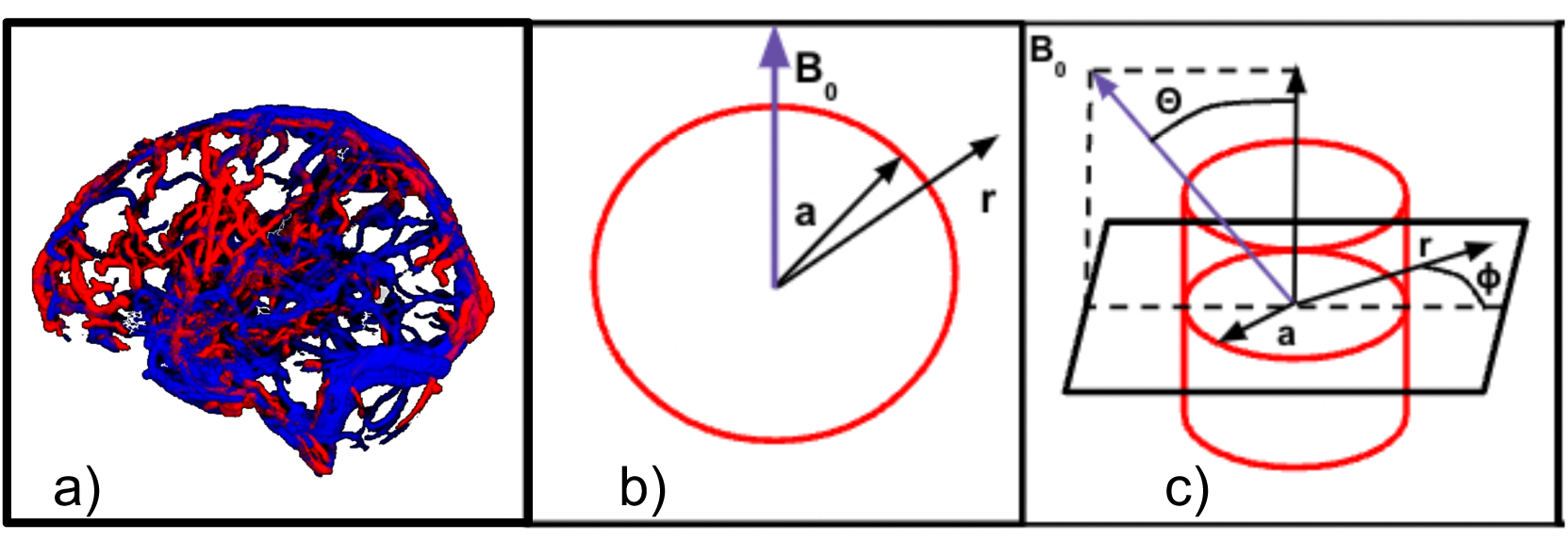
Illustration of three vascular models used to predict the rs-fMRI BOLD signal. a) vascular-based mVAN simulation (mVAN Model), b) the two-dimensional (2D) infinite cylinder model (2D Cylinder Model), c) the three-dimensional (3D) infinite cylinder model (3D Cylinder Model). Here, *a* is the cylinder radius, *r* is the distance between the point of interest and the centre of the cylinder cross-section, B_0_ is the main magnetic field, θ is the angle between the B_0_ and the cylinder axis, and φ is the angle between the vector <**r**> and the projection of B_0_ on the plane perpendicular to the cylinder axis.

**Table 1.**
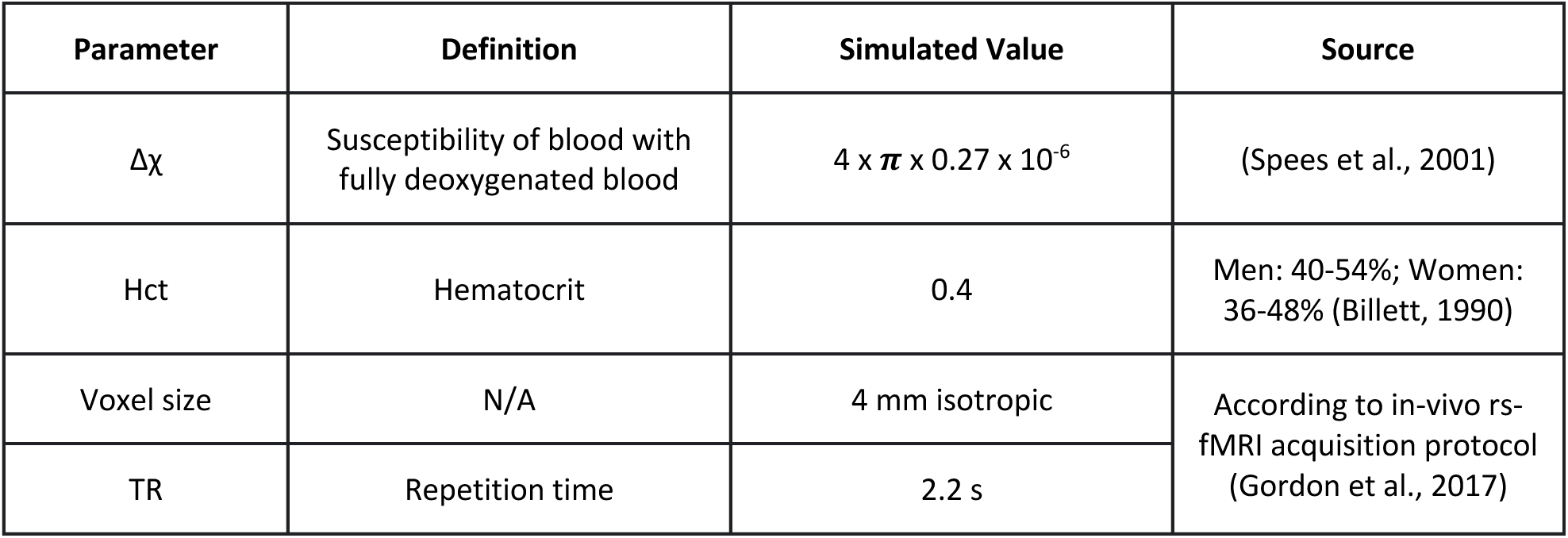
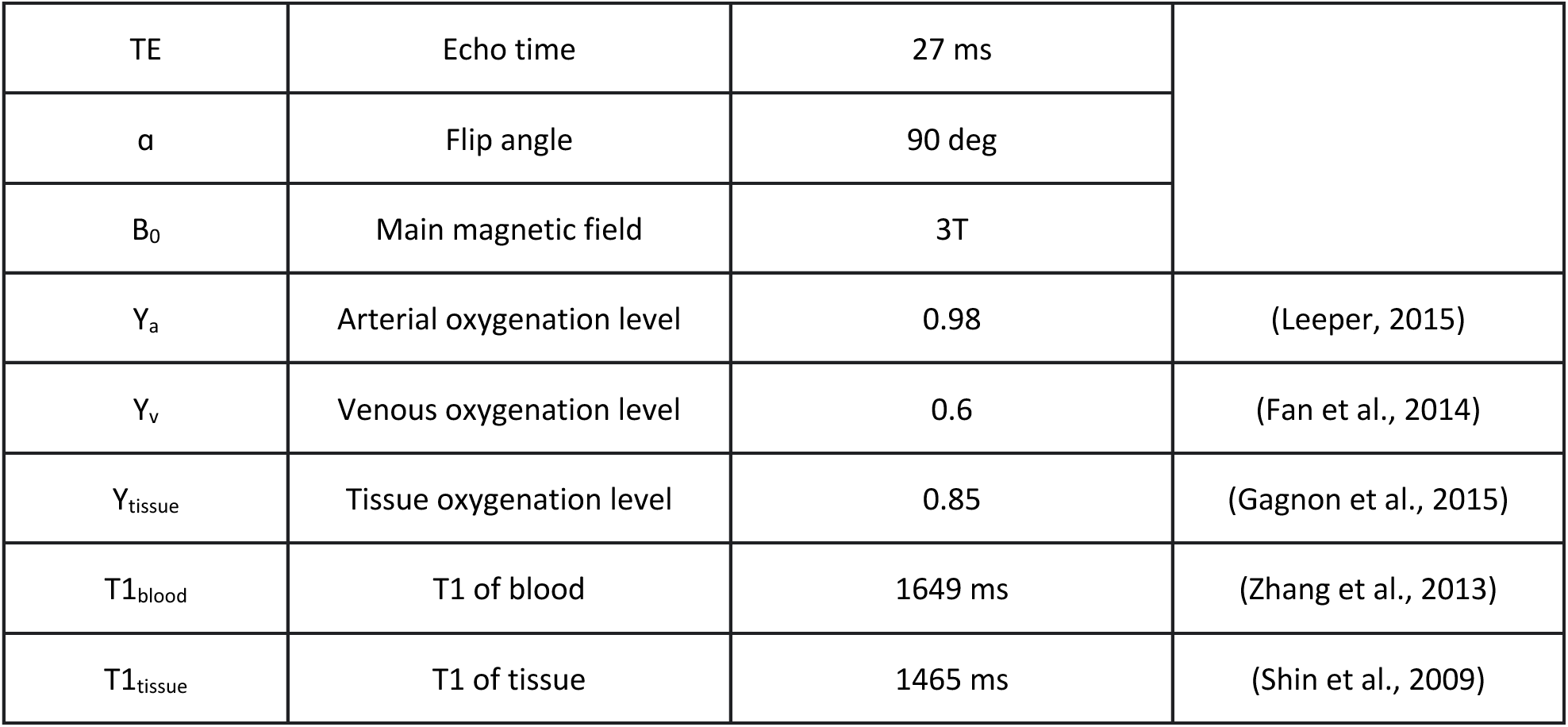
Simulation parameters and values. For all three simulation models, these values were set to default unless otherwise stated.

#### 2.5.3 mVAN Model: 3D macrovascular mVAN

In this model, magnetic susceptibility maps are based on experimentally acquired mVANs (**Fig. 3**). A whole-brain susceptibility map was generated numerically using the Fourier method, whereby a mask of the vasculature was generated by upsampling the mVANs to 0.2 mm isotropic resolution, and necessary zero-padding by the size of a full field-of-view on each side of the matrix to avoid cycle convolution wraparound. The experimental data and simulated data were first aligned with the coordinates extracted from the processed macrovascular segmentation, as described earlier. To minimise computational complexity, the susceptibility map was derived using a Fourier-based model-independent convolution approach (**Fig. 3**) (Eq. 1-2) (Cheng et al., 2009; Salomir et al., 2003), as follows.

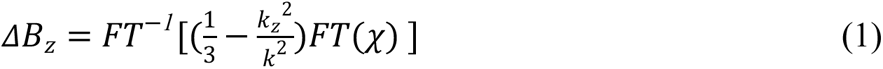

**Figure 3.**
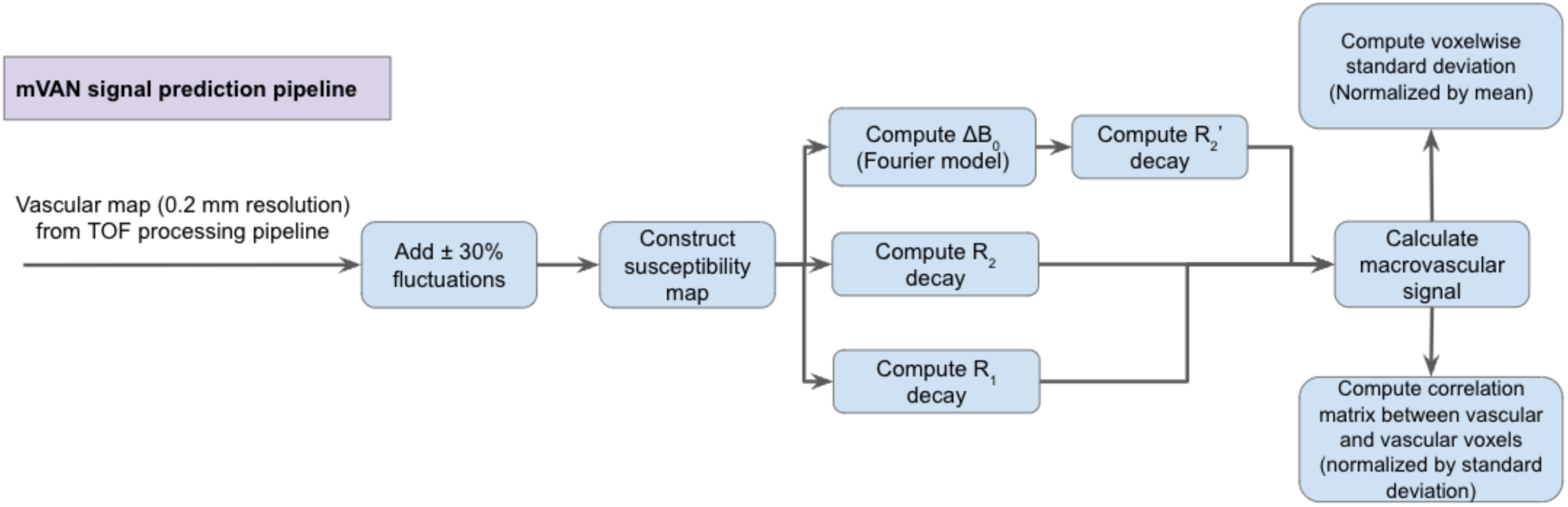
Overview of the simulation procedure (based on vascular segmentation).

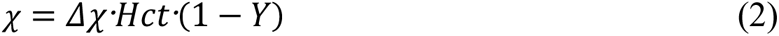

FT denotes the Fourier transform, χ the local susceptibility, *k*_z_ the distance in the k space along the z-axis and *k* the distance in k-space (*k*^2^ = *k*_x_^2^ + *k*_y_^2^ + *k*_z_^2^). R_2_^’^ is then calculated through the magnitude of the complex-valued mean magnetization of the dephasing spins resulting from the B_0_ offset. We added the necessary zero-padding by the size of a full field-of-view on each side of the matrix to avoid cyclic convolution wraparound.

The experimental data and simulated data were first aligned with the coordinates extracted from the processed macrovascular segmentation, as described above. Then, to resample the phase offset map to fMRI spatial resolution, the phase offset in mVAN Model was summed across 8000 neighbouring 0.2 mm voxels (which sum up to a 4 mm isotropic voxel) to calculate the total BOLD signal. We further assumed the tissue type outside macrovasculature was grey matter (GM) to calculate the appropriate susceptibility difference between blood and tissue. The mean BOLD signal was calculated as defined by Eq. 3 and 4 as

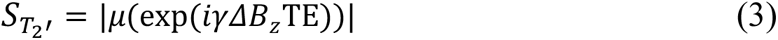

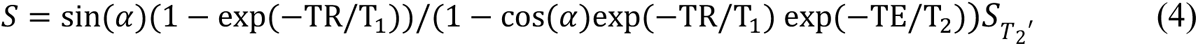

where γ is the gyromagnetic ratio, and the operator |μ(⋅)| represents the magnitude of the mean of a complex number.

#### 2.5.1 2D Cylinder Model: 2D infinite cylinder

On the other end of the spectrum of complexity, we tested a single-voxel two-dimensional (2D) infinite-cylinder model, in which field offset is estimated in an analytical approach (**Fig. 4a**) (Eq. 5 and 6) (Ogawa, Menon, et al., 1993). Here, instead of generating a volumetric mask of the mVAN, we calculated at each location of the mVAN the diameter and orientation of the best-fitting cylinder and used this information (combined with the blood-tissue susceptibility difference) to calculate analytically the magnetic field offset. This calculation is based on a closed-form expression for the extravascular and intravascular field offsets that are a function of the vessel geometry and blood susceptibility, i.e.,

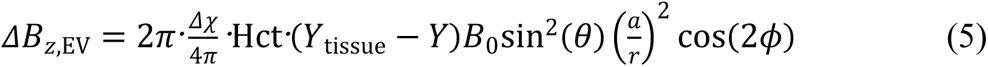

**Figure 4.**
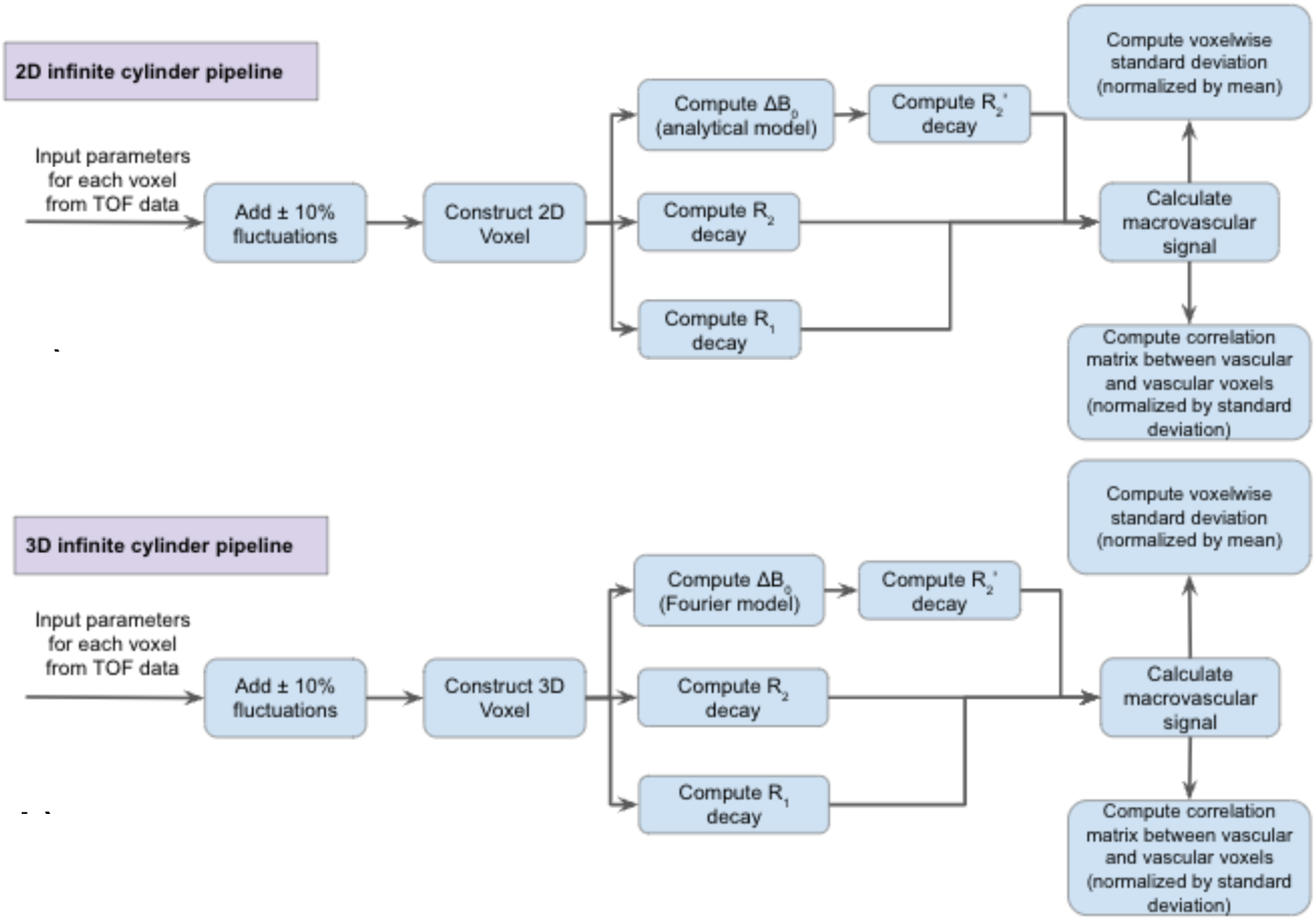
Overview of the simulation procedures for Models 2 and 3 (infinite-cylinder based). To predict the rs-fMRI signal, three models were used, namely a) the two-dimensional (2D) infinite cylinder model, b) the three-dimensional (3D) infinite cylinder model. The simulation parameters were extracted from the TOF data.

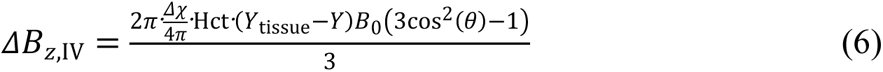

where ω_0_ is the main magnetic field in terms of angular frequency, Δχ: magnetic susceptibility of fully deoxygenated blood, Hct: Hematocrit, *Y* is the percent blood oxygenation (*Y*_a_ for artery and *Y*_v_ for vein), (Y_tissue_−Y) is the degree of deoxygenation difference between blood and tissue, θ is the angle between the B_0_ and the vascular axis, *a* is the cylinder radius, *r* is the distance between the point of interest and the centre of vascular cross-section, φ is the angle between vector <**r**> and B_0_ projection in the plane.

This model assumes that the shape of the blood vessel is an infinite cylinder with zenith θ, azimuth ɸ, and radius *a* derived from fBV. These input parameters can all be obtained from the arterial and venous segmentations based on the TOF data. This 2D modelling is suitable when it is assumed that the magnetic susceptibility does not vary along the length of the infinite cylinder, and facilitates our understanding of the dependence of IV and EV rs-fMRI signals on a single large vessel. As in the case in the work of Bandettini and Wong (Bandettini & Wong, 1995), we constructed a 2D voxel for each vascular orientation-fBV combination derived from the experimental data, divided into 4,000 × 4,000 1-μm isotropic sub-voxels, with each voxel defined by a set of vascular (arterial or venous) orientation and fBV values derived from the TOF data in the previous step. The 2D model is more conducive to analytical calculations of susceptibility, and the dimensionality reduction can be traded for higher spatial resolution.

#### 2.5.2 3D Cylinder Model: 3D infinite cylinder

This was a 3D version of the infinite cylinder model (also assumed in 2D Cylinder Model), with the exception that the vessel is not assumed to penetrate the voxel at a right angle, such that fBV will vary with θ, and that the susceptibility pattern can vary along the length of the vessel. We constructed a 3D voxel containing 100×100×100 40-μm isotropic sub-voxels, in which the infinite cylinder is defined by the collection of measured parameters as was done in the 2D Cylinder Model Model. The field-offset estimation procedure is the same numerical Fourier-transform-based method as used in the mVAN Model. This model is an intermediate between the mVAN Model and the 2D Cylinder Model, and is evaluated as a less computationally intensive approximation of the mVAN Model. The 2D and 3D Cylinder Model are both single-voxel and thus do not take into account interactions between neighbouring vessels that may influence EV field offsets.

#### 2.5.4 Temporal dynamics of the simulated vasculature

Vascular dynamics were simulated as sinusoidal variations at 0.1 Hz (Chu et al., 2018) for arterial fBV and venous *Y* (Uludağ & Blinder, 2018; Vazquez et al., 2013). We used a sinusoid with ±10% variations (peak-to-peak) for Y and fBV fluctuations in the 2D and 3D CylinderModels. For mVAN Model, the mVAN was upsampled to 0.2 mm isotropic resolution, and the smallest detectable vessels had approximate diameters of 0.8 mm. Due to the coarser sample, the circular cross-section of each 0.8 mm vessel, assuming it is at the centre of its voxel, would be upsampled in 2D into a 13-voxel neighbourhood of 0.2-mm isotropic voxels. Thus, it was necessary to expand the fBV variations to ±30% to allow the quantized vascular fBV to fluctuate by one voxel in the diameter dimension. The simulated BOLD signal can be calculated as Eq. 4.

From these simulated BOLD signals, we computed both resting-state fluctuation amplitudes (RSFA) and FC (i.e., the correlation between voxel time courses). For computing the RSFA, both the simulated and experimental BOLD signals were normalized by their respective means, with outlier time points (defined as lying beyond 1.5 quartiles of the mean) removed. Simulated FC metrics were calculated for voxelwise arterial-arterial, venous-venous, and arterial-venous correlations with the simulated signals from all three models. As the simulated data is noiseless but is to be compared with noisy experimental results, we added noise for additional realism. Thus, as suggested by Golestani et al. (Golestani & Goodyear, 2011), we used the temporal standard deviations of simulated time courses as a surrogate for SNR; that is, with noise being similar across voxels, a higher baseline signal fluctuation would translate into a higher SNR. We then multiplied the cross-correlation coefficients in the FC metrics with the σ for each simulated voxel to obtain SNR-independent FC measurements.

The simulations were conducted using our servers equipped with 14 cores of Intel Xeon X5687 CPU (at 3.6 GHz) (Intel Corporation, Santa Clara, CA, United States) and 180 GB of memory running Red Hat Enterprise Linux Server 7.7 (Red Hat Inc., Raleigh, NC, United States). A customised simulation script was written in MatLab 2019b (MathWorks Inc., Natick, MA, United States.).

#### 2.5.5 Perivascular signal contributions

Predicted (mVAN Model only) and measured rs-fMRI signals were compared in the perivascular tissue as well. **Figure 1c** illustrates the methods used to assess perivascular effects of the macrovasculature on the rs-fMRI signal, whereby the perivascular ROIs were generated as follows. First, the arterial and venous segmentations were dilated by 1 voxel in 3D. Second, the original vascular ROIs were subtracted from the dilated vascular ROIs. This produced the extravascular ROI that is centred at 1 voxel distant from the centre of the vasculature. This process was repeated for perivascular distances of 2 and 3 voxels. With the approaches discussed in the previous section, cross-correlation-based connectivity values were calculated separately for BOLD signals in arterial and peri-arterial ROIs, and between venous and peri-venous ROIs. This was done for both simulated and in-vivo experimental signals.

### 2.6 Agreement between predicted and measured correlation coefficients

Each simulated voxel will result in the simulated macrovascular-based BOLD signal corresponding to a vascular location in the experimental data, which corresponds to measured fluctuations of the BOLD data from the same voxel. In this way, to quantify model prediction accuracy, the simulated RSFA and vascular-driven correlation coefficients predicted by all three models were regressed against the experimental cross-correlation coefficients. All vascular-related correlation coefficients (simulated and experimental), which were used as measures of FC, were binned. Given that there are more voxels for venous ROIs than arterial ROIs), the bin sizes were adapted approximately proportionately, i.e., 100 for arterial-arterial correlations, 1000 for arterial-venous correlations, and 10000 for venous-venous correlations (as there are more venous voxels than arterial voxels), to ensure a similar number of fitted data sets for all three categories. This was done separately for each dataset. The R^2^ values were calculated to assess the quality of the prediction of measured by the simulated FC values. To determine how well the simulated model can account for perivascular effects, a similar regression procedure was conducted for the perivascular FC metrics using mVAN Model. The correlations were grouped by distance from the edge of the vasculature (i.e., 1, 2 or 3 voxels away). Similarly, the procedure described above was applied to regressing simulated and against experimental perivascular BOLD signal fluctuation amplitudes, separately for each vascular distance.

## 3 Results

### 3.1 Theoretical predictions based on a 2D infinite cylinder

The analytical equations (Eq. 5, 6) allowed us to first gain an intuitive understanding of the interactions between the BOLD signal and various vascular parameters. When simulated based entirely on the default parameters listed in Table 1, arterial R_2_’ increased with fBV but only up to fBV =< 0.4, and decreased as fBV increased beyond 0.4 (**Fig. 5a**). This is consistent with decreasing phase coherence as fBV increases in a tissue-dominated voxel followed by an increasing phase coherence once the voxel became blood-dominated. However, as the vascular orientation varied, so did the behaviour of R_2_’ (**Fig. 5b**). The peak R_2_’ was generally not found at the θ extrema (corresponding to when the vessel is parallel or perpendicular to B_0_). On the venous side, in this case, R_2_’ declined with increasing Y_v_, but this behaviour also varied with fBV. Venous R_2_’ was maximized at an intermediate value of fBV. As shown in **Fig. 5c**, it was higher for fBV = 0.4 than for lower and higher fBVs (0.1 and 0.7, respectively). Lastly, for the default fBV, like arterial R_2_’, the venous R_2_’ dependence on Y_v_ varied with θ, with R_2_’ higher at an intermediate value of 3π/4 than at θ of π/2 or π(**Fig. 5d**). BOLD signal intensity was disproportional to R_2_’ (**Fig. 5e-h**): the R_2_’ was maximized at intermediate values of fBV, the baseline BOLD signal intensity was minimized at intermediate values of fBV; when the R_2_’ was maximized at intermediate theta values, the baseline BOLD signal intensity minimized at intermediate theta values.

**Figure 5.**
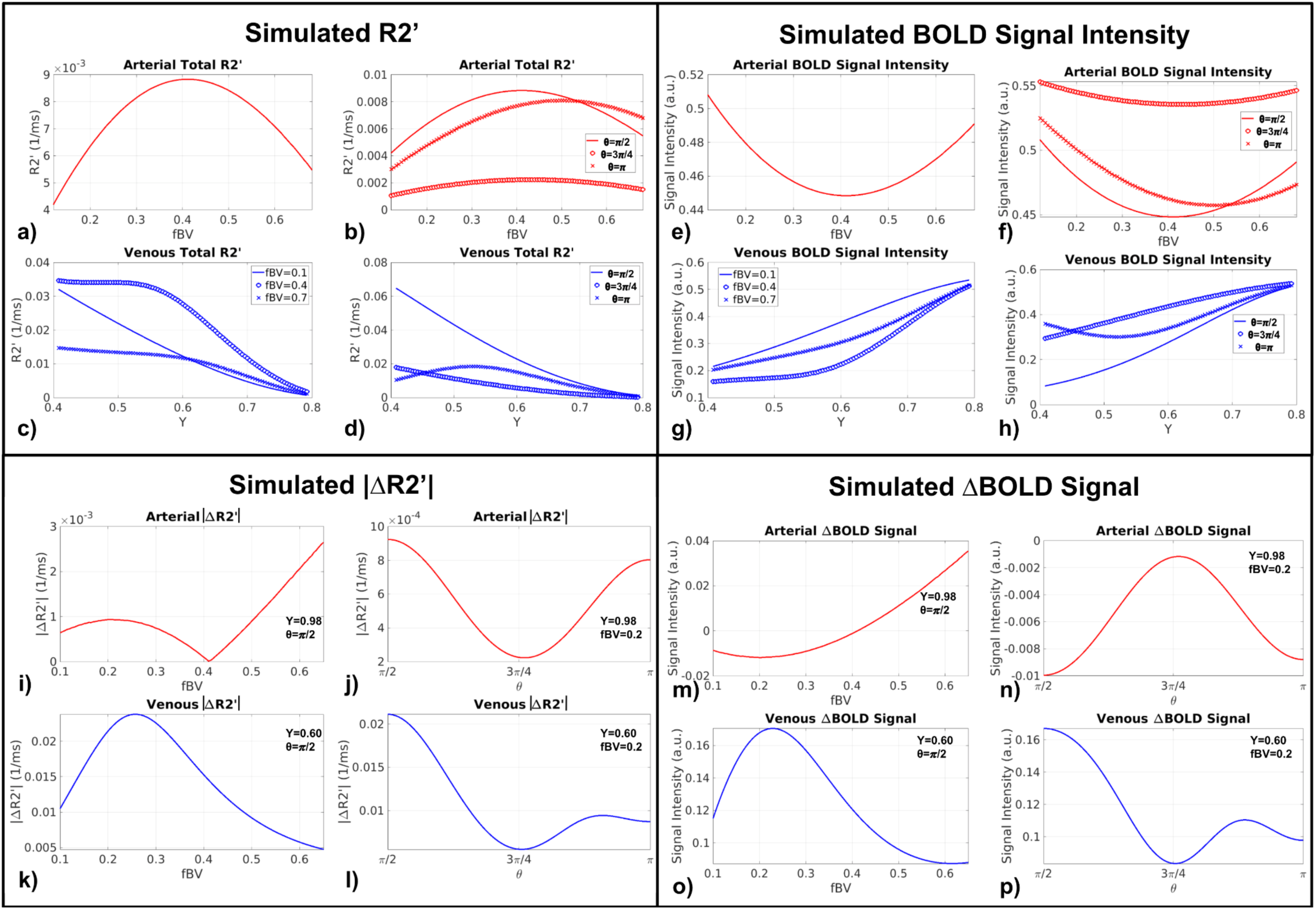
Influence of macrovascular fBV and *Y* on R_2_’ and BOLD signal intensity, simulated analytically using the 2D Cylinder Model (a-h) and influence of macrovascular θ and fBV on resting-state R_2_’ fluctuations (|ΔR_2_’|) and BOLD fluctuation (ΔBOLD) (i-p). All simulations were performed using the default simulation parameters listed in Table 1, and assuming 10% peak-to-peak oscillations in fBV (for arteries) or Y_v_ (for veins).

In rs-fMRI, fluctuations in R_2_’, rather than R_2_’ itself, is more relevant. The quantity ΔR_2_’, computed as the difference between the maximum and minimum R_2_’ (associated with the maxima and minima in Y and fBV), can be viewed as the derivative of R_2_’ relative to the oscillations in Y and fBV (**Fig. 5a-d**). Arterial ΔR_2_’ initially rose with increasing fBV, then decreased as fBV surpassed ∼0.2, eventually increasing again for fBV > 0.4 (**Fig. 5i**). Arterial ΔR2’ decreased with θ as θ increases to 3π/4 and then increased for θ up to π (minimum at θ = 3π/4) (**Fig. 5j**). Venous ΔR_2_’ in (**Fig. 5k**) showed the opposite trend as arterial with fBV < 0.4 (**Fig. 5i**), decreasing with increasing fBV until fBV ∼0.2, then increasing for higher fBVs. Lastly, like arterial |ΔR_2_’|, venous |ΔR_2_’| also decreased with θ until θ = 3π/4, then increased for greater values of θ. Moreover, ΔR_2_’ became more stable while θ approached π. ΔBOLD was in turn computed as the difference in the BOLD signal corresponding to the two extremes of the input signals (fBV for artery and Y for vein), as derived from the associated R_2_’ values. As seen in **Fig. 5m-p**, the magnitude of ΔBOLD can be viewed as proportional to |ΔR_2_’|, whereas the polarity of ΔBOLD will influence the correlation between ΔBOLD time courses. For instance, the venous ΔBOLD in (**Fig. 5o**) showed the opposite trend as arterial ΔBOLD (**Fig. 5m**), decreasing with increasing fBV until fBV ∼0.2, increasing for higher fBVs and remaining positive throughout all fBVs. Thus, an equivalent increase in fBV for both an artery and a vein would lead to their BOLD time courses remaining negatively correlated. Also, arterial and venous ΔBOLD showed the same magnitude variation trend until arterial ΔBOLD changed polarity at fBV ∼0.4, which suggests a negative correlation between arterial and venous BOLD time courses for fBV > 0.4. Lastly, arterial ΔBOLD increased with θ until θ = 3π/4, then increased for greater values of θ. This is contrary to the trend observed for venous ΔBOLD (peaking at θ = 3π/4) (**Fig. 5p**).

### 3.2 Comparison between simulated and measured BOLD RSFA

As shown in **Fig. 6a and d** for a representative data set, arterial RSFA is more poorly predicted by the mVAN Model than venous RSFA. For the 2D Cylinder Model, but the goodness of fit was limited for both arterial and venous RSFA (**Fig. 6b, e**). In both cases, the low goodness of fit may be attributed to a large cluster of measured BOLD RSFA values in voxels that exhibit a low simulated RSFA. The 3D Cylinder Model is an improvement on the 2D Cylinder Model especially for predicting venous RSFA, but the performance remains lower than for the mVAN Model (**Fig. 6c, f**). These trends are generalizable to all subjects (see *Supplementary Materials*). See **Fig. S2** for results for all subjects.

**Figure 6.**
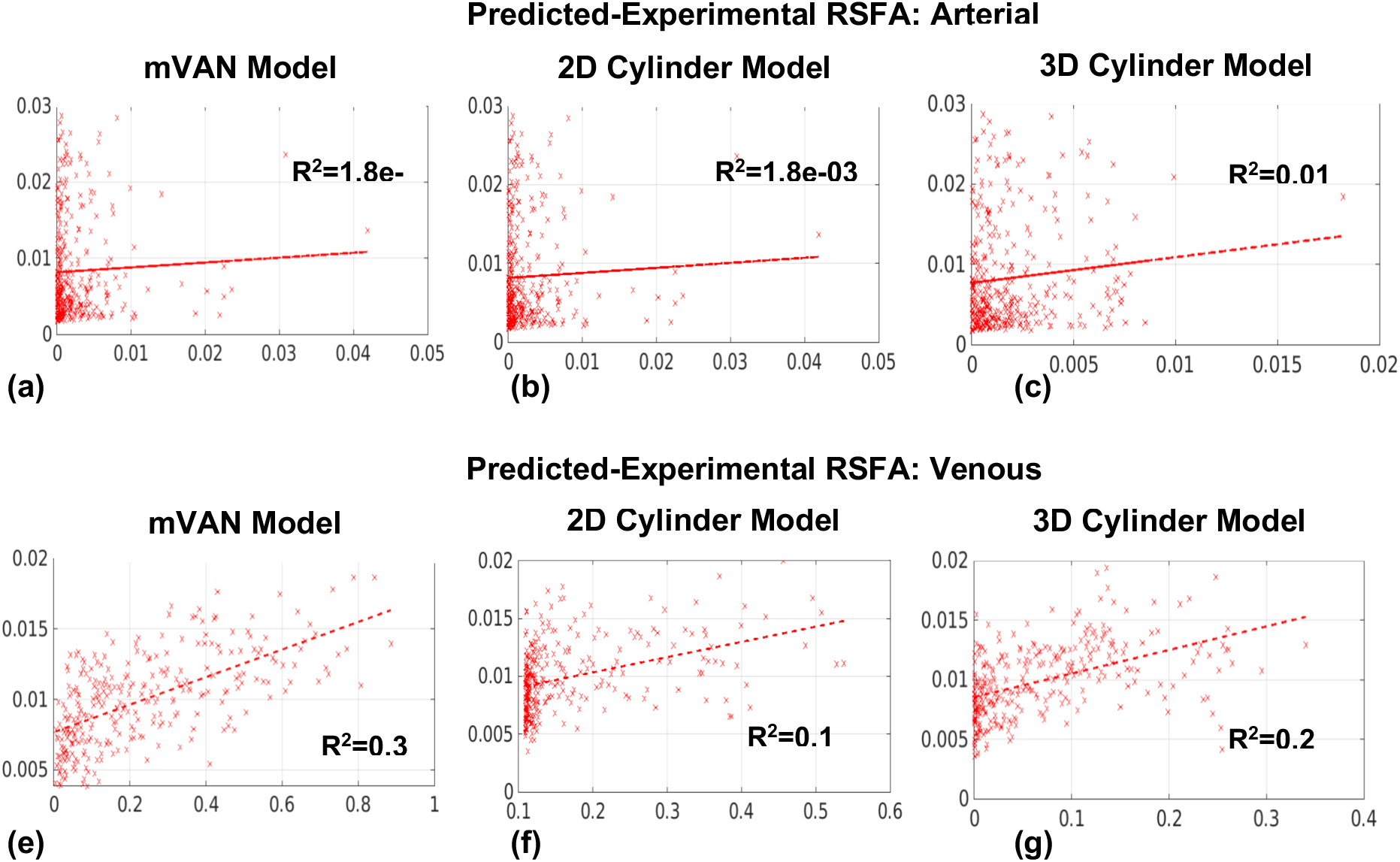
Regression between the in-vivo predicted and experimental vascular RSFAs. Regressions are shown between the in-vivo experimental and simulated RSFA for mVAN Model (a,b), 2D Cylinder Model (c,d) and 3D Cylinder Model (e,f). Data are from a representative subject. Each dot represents a bin average.

### 3.3 Comparison between predicted and measured correlation coefficients

The simulated correlation coefficients based on the mVAN Model show the highest predictiveness for venous-venous correlation coefficients (**Fig. 7g**), followed by arterial-venous correlation coefficients (**Fig. 7d**), and with the lowest predictability, arterial-arterial correlation coefficients (**Fig. 7a**). For the 3D Cylinder Model, as expected, while most regressions showed significant positive associations between simulated and in-vivo correlation coefficients, the goodness of fit was low (**Fig. 7b, e, h)**, with the arterial-arterial correlation R^2^ being the lowest (< 0.01). We also noted that the data points appear to subdivide into two groups, one spanning correlation coefficients of > 0.04, and the other one spanning the area between 0 and 0.04. As our regression does not make provisions for these two regimes of correlations. For the 2D Cylinder Model, the predictability of venous-venous correlations remains remarkably high (**Fig. 7i)**, followed by arterial-venous correlations (**Fig. 7f**). The accuracy of the prediction of arterial-arterial correlations was still relatively low (**Fig. 7e**). See **Figs. S3-S5** for results for all subjects.

**Figure 7.**
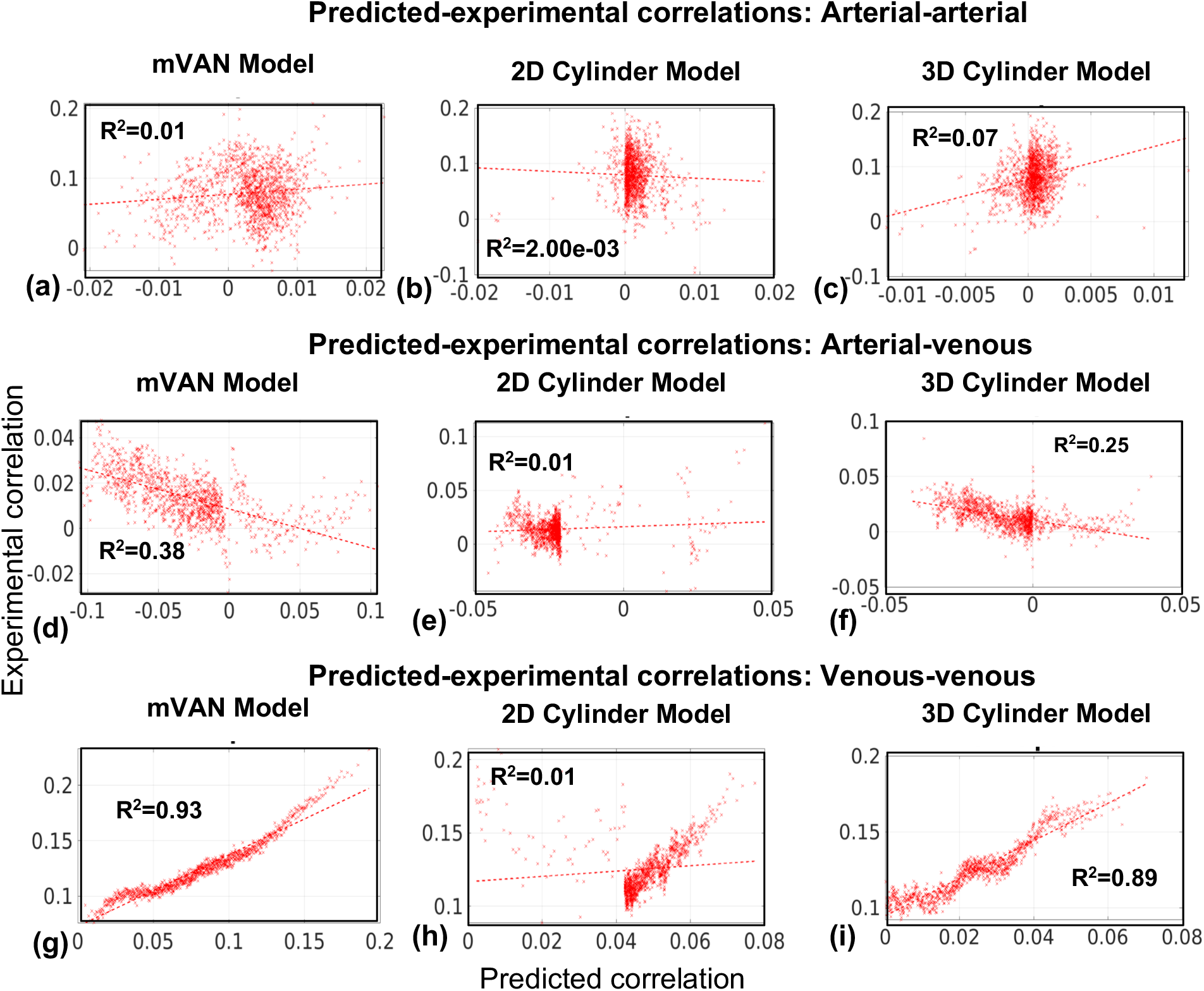
Regression between the in-vivo predicted and experimental vascular correlation coefficients. Data is from the same subject as in Fig. 6. Regression lines for arterial-arterial correlations,arterial-venous correlation, and venous-venous correlation coefficients are shown for all models. Each dot represents a bin average.

### 3.4 Perivascular effects

#### 3.4.1 Perivascular contributions to GM BOLD signal standard deviations: predictivity using simulations

Only the mVAN Model provided the possibility to examine the perivascular effects. We found the majority of vascular contributions to the surrounding tissue to be limited to one voxel away from the original vascular masks (**Fig. S6** in Supplementary Materials). As seen in **Fig. 8**, both arterial and venous perivascular RSFA are poorly predicted by the mVAN Model, irrespective of the distance from the vascular voxels. The predicted RSFA values cannot account for the large spread of experimental RSFA values.

**Figure 8.**
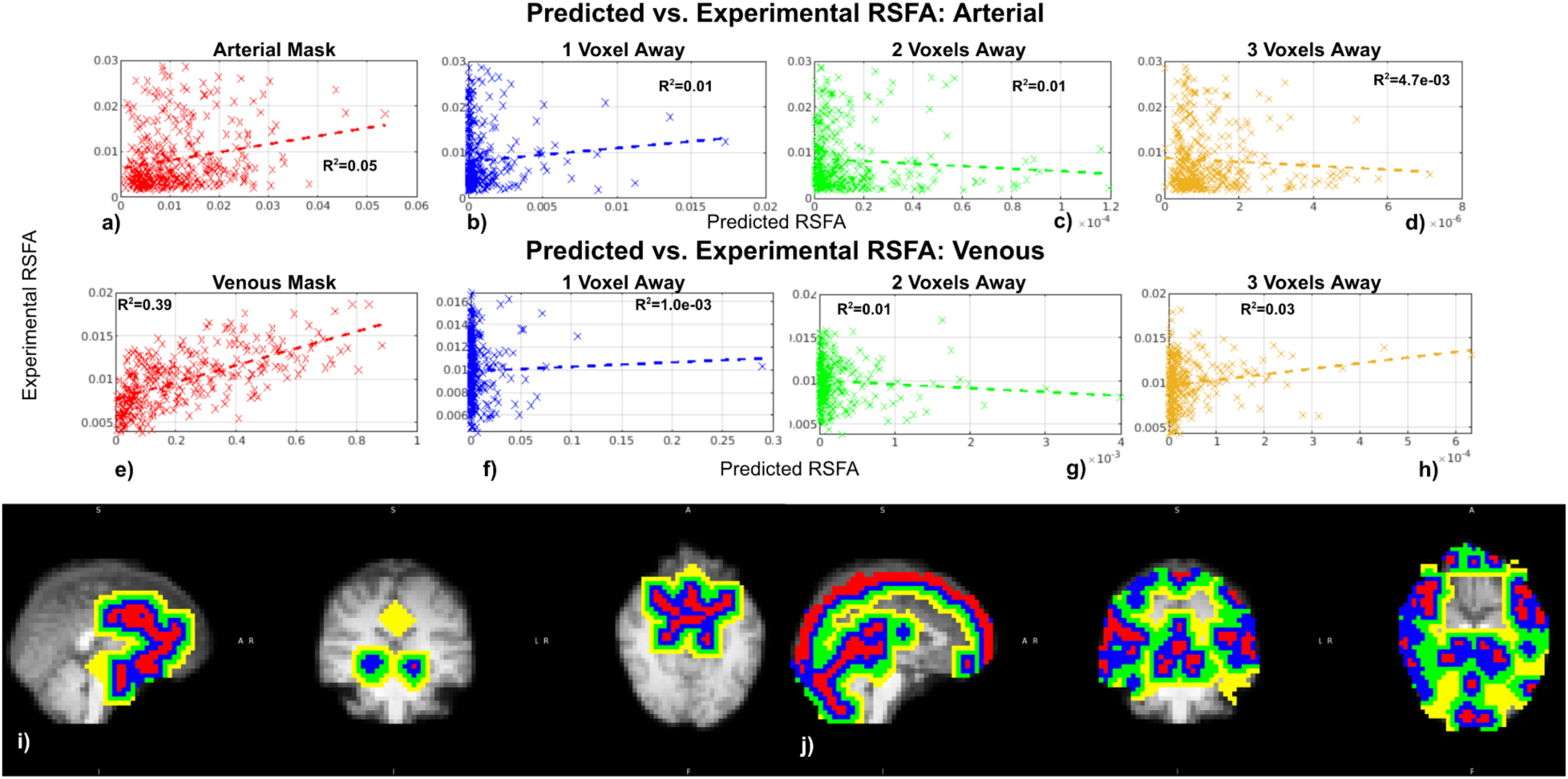
Dependence of perivascular BOLD standard deviation on distance from vasculature: predicted vs. experimental results (the mVAN Model). Data taken from the same representative subject as in Fig. 6. Perivascular masks are shown for both arterial and venous, with colour coding as: red – intravascular, blue – one voxel away from the vascular voxel, green – two voxels away, and yellow – three voxels away (i,j).

#### 3.4.3 Perivascular contributions to GM BOLD signal correlations: predictiveness using simulations

Unlike for venous RSFA, venous-driven correlation remains quite well predicted by the mVAN Model at 1 voxel away from the vascular voxel, as shown by the R^2^ value (**Fig. 9f**). The regression remains significant at two and three voxels away, however R^2^ decreased substantially (**Fig. 9g, h**). The predictability of arterial-driven correlations remains poor for all perivascular distances (**Fig. 9a-d**). See **Figs. S7-S8** for results for all subjects.

**Figure 9.**
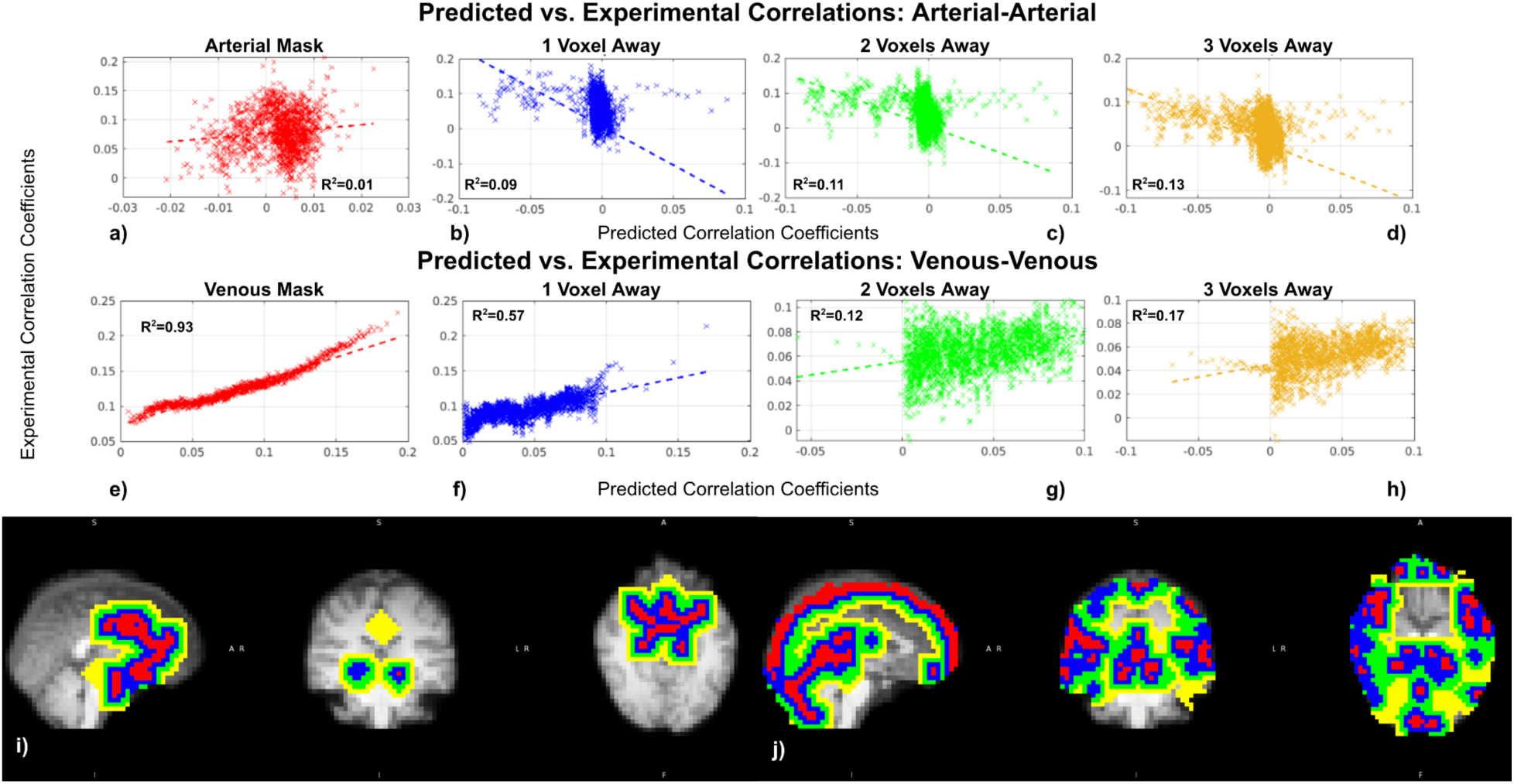
Dependence of perivascular correlation coefficients on distance from vasculature: predicted vs. experimental results (mVAN Model). Data were taken from the same representative subject as in Fig. 6. An illustration of arterial-arterial correlation (a-d) and venous-venous correlation (e-h). Perivascular masks are shown for both arterial and venous, with colour coding as: red – intravascular, blue – one voxel away from the vascular voxel, green – two voxels away, and yellow – three voxels away (i,j).

### 3.5 Comparison of three models across all subjects

Comparisons of R^2^ values from the three models against measurements from all 4 subjects are summarised in **Figure 10**. In general, the mVAN Model provided the highest R^2^ values, followed by the 3D Cylinder Model, and lastly the 2D Cylinder Model (**Fig. 10a-c)**. Additionally, the simulated venous-venous correlation coefficients (**Fig. 10a**) are associated with the highest R^2^ values (all < 0.8), followed by arterial-venous correlations (**Fig. 10b**), and then arterial-arterial correlations (**Fig. 10c**). Perivascular prediction differences are also summarised in **Fig. 10** across all subjects and distances. Generally, the R^2^ values of arterial-arterial correlations were low, and increased with distance away from the original vascular mask (**Fig. 10e**). As with the correlation coefficient simulation, the RSFA simulation shows a similar pattern, but with generally lower R^2^ than for correlation coefficients. Both arterial and venous signals were better fitted by the mVAN Model than by the other models (**Fig. 10f,g**). Moreover, there is a trend of decreasing goodness of fit as the distance from macrovasculature increases (**Fig. 10h,i**).

**Figure 10.**
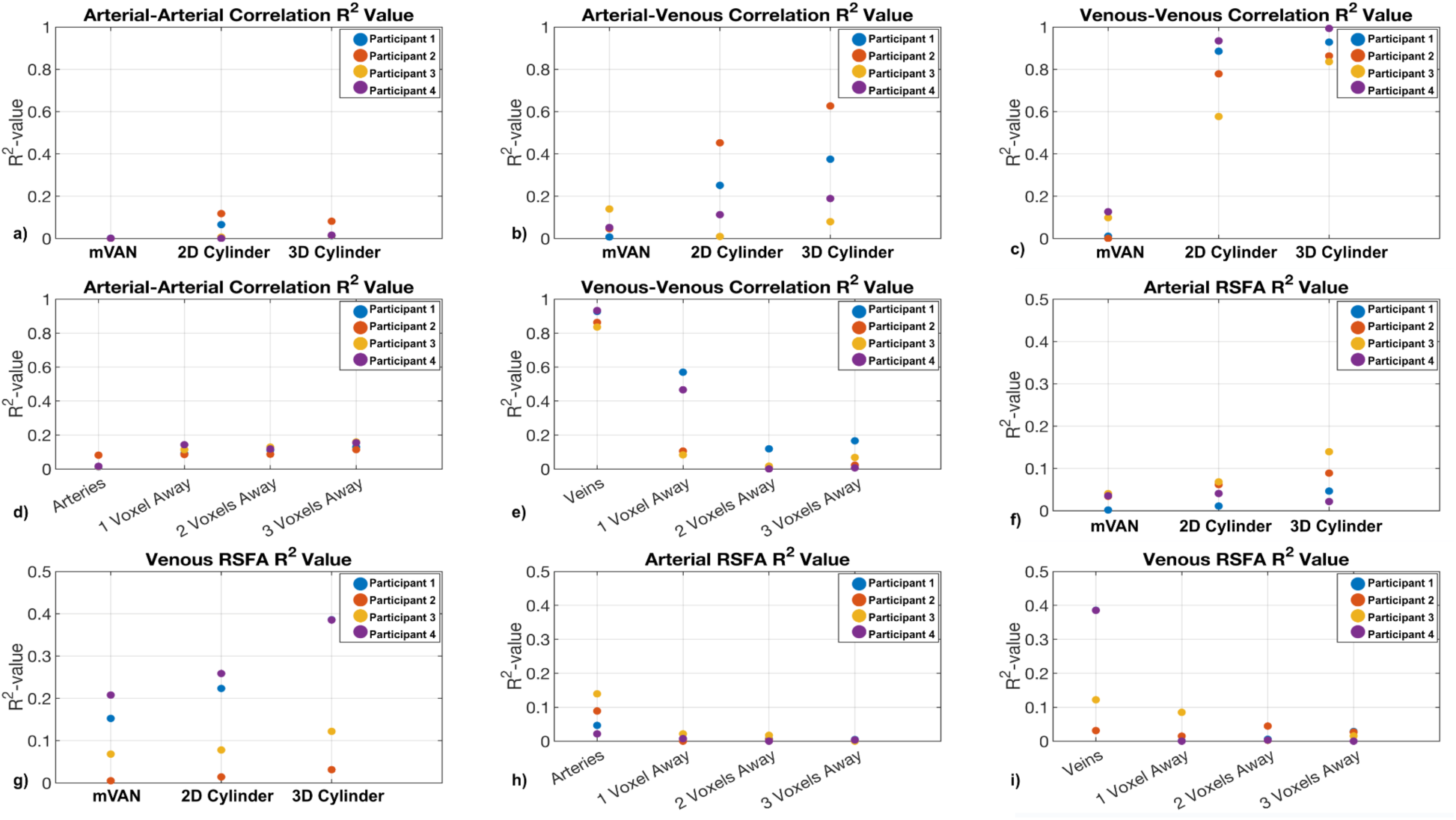
Summary of results from all subjects: the agreement of predicted and experimental vascular FC in terms of correlations, quantified by R^2^. R^2^ values are plotted for all 4 data sets for (a) arterial-arterial correlation, (b) arterial-venous correlation, (c) venous-venous correlation, (d) arterial-arterial perivascular correlation, (e) venous-venous perivascular correlation, (f) arterial standard deviation, (g) venous standard deviation, (h) arterial perivascular standard deviation, and (i) venous perivascular standard deviation.

## 4 Discussion

Despite the strong influence of macrovascular contributions to BOLD fMRI (Huck et al., 2023; Menon, 2012; Tong et al., 2015; Zhong et al., 2023), previous research demonstrated that removing this bias is not an easy task (Huck et al., 2023; Menon, 2002; Ragot & Chen, 2019; Stanley et al., 2021). Using analytical simulations informed by experimental data, the present study examines biophysical-simulation-based approaches to predict the macrovascular BOLD contributions proximal and distal to the vasculature. We explored the use of a full mVAN model as well as an infinite-cylinder approximation (2D and 3D). We found that:

1. With the availability of angiographic data, it is feasible to model macrovascular BOLD FC using both the mVAN-based model and 3D infinite cylinder models, though the former performed better.
2. Biophysical modelling can accurately predict the BOLD FC (in terms of correlations) for large veins, but not for large arteries.
3. Biophysical modelling provided less accurate predictions for RSFA than for FC.
4. Modelling of perivascular BOLD connectivity was feasible at distances close to veins but not arteries, with performance deteriorating with increasing distance.

### 4.1 Origins of the resting-state macrovascular BOLD fluctuations

The macrovascular contribution to the rs-fMRI BOLD signal has many physiological origins. The concept of systematic low-frequency oscillations (sLFOs) was first proposed in an earlier review article and defined as a low-frequency vasogenic BOLD signal travelling through the brain (Tong et al., 2015). This phenomenon does not appear to originate within the brain (Frederick & Tong, 2010; Li et al., 2018; Tong & Frederick, 2010) and is likely associated with heart rate variability (Thayer et al., 2012), gastric oscillations (Mohamed Yacin et al., 2011; Rebollo et al., 2018), vasomotion (Hundley et al., 1988; Mayhew et al., 1996; Rivadulla et al., 2011), respiratory volume variability (Birn et al., 2006; Chang et al., 2009) and/or variations in carbon dioxide levels (Sassaroli et al., 2012; Wise et al., 2004). In previous studies using near-infrared spectroscopy, it has also been demonstrated that such oscillations travel through the vasculature. This can also be demonstrated in the regression of finger-tip oxygenation time course with whole-brain BOLD signals in a spatially coherent manner, as demonstrated in several previous studies (Frederick et al., 2013; Tong, Bergethon, et al., 2011; Tong et al., 2014, 2015; Tong, Hocke, et al., 2011; Tong, Lindsey, et al., 2011; Tong & Frederick, 2012, 2014). As venous blood is predominantly responsible for the BOLD signal (Ogawa, Lee, et al., 1993; Segebarth et al., 1994), the presence of sLFOs may suggest that certain findings from rs-fMRI experiments may not reflect neural activity but rather venous bias (Aso et al., 2020).

To understand the influence of the macrovasculature, it is essential to have a basic biophysical understanding of the macrovascular BOLD effect. The macrovascular contribution is likely to be strongest within the confines of both macrovascular and perivascular tissue, as discussed in our previous study (Zhong et al., 2023). The origins of the large variability in the perivascular BOLD effect across brain regions and acquisitions (Huck et al., 2023; Zhong et al., 2023) are unclear from in-vivo experiments alone. However, we hope that our simulated results can contribute to a better understanding of this phenomenon.

### 4.2 Comparison of simulation models

#### 4.2.1 Theoretical comparisons of our models

Three simulation models were compared in this study, ranging in complexity and realism from low to high, namely from the 2D infinite-cylinder model, to the 3D infinite-cylinder model, to the empirical mVAN model. The 3D Cylinder Model is expected to be more accurate than the 2D Cylinder Model due to its ability to capture a geometric effect of orientations (azimuth and zenith) on fBV. That is, the 2D Cylinder Model assumes that the effect of vessel orientation on fBV is negligible, but unlike microvascular BOLD simulations (Berman et al., 2018), macrovascular simulations cannot ignore fBV changes caused by vessel obliquity, as macrovascular fBV is more substantial and arises from much larger “cylinders”. As an example, a vessel with an azimuth angle of 45° or zenith angle of 45° would have a fBV of approximately 1.41 times that of a vessel with both angles at 0. Thus, the 2D Cylinder Model is expected to be outperformed by the 3D Cylinder Model, which also models vessels as cylinders but more accurately captures fBV and vessel orientation. While the 2D Cylinder Model was expected to provide low accuracy, it was unclear a priori how these simplifying assumptions would affect our BOLD predictions, and given its practical advantages it was evaluated in our study. Our conclusion is that, for modelling large vessels, this model is inadequate and the disadvantages outweigh the advantages.

Likewise, we expected the mVAN Model to produce the most realistic BOLD simulations since it incorporates the shape and position effects of real macrovasculature that cannot be captured by the 2D and 3D Cylinder Models. The mVAN Model would be able to capture details missing from the 3D Cylinder Model, such as the geometryof blood vessels (with bends, branching, etc), partial volume effects, and positional effects (when the vessel is not centred in a voxel). Despite the advantages of the mVAN Model, it still has some limitations. Because it is based on the Fourier-based discrete Green’s function, the accuracy of this approach depends on the grid resolution. More specifically, for high accuracy, the diameter of the smallest blood vessels should span at least four voxels (Tang et al., 1993). Coarse spatial sampling of the vascular susceptibility gradient and boundaries leads to truncation or “ringing” effects, which can be alleviated by spatially upsampling, thus smoothing the kernel function. Conversely, due to its simpler geometry, the 2D Cylinder Mode can be applied at arbitrarily high spatial resolution (by using smaller sub-voxels) to provide a more accurate representation of fBV variations, which is especially relevant for small fluctuations in the resting state. Furthermore, given the complexity of the mVAN Model, a more intuitive understanding of the BOLD signal dependence on vascular geometry is challenging. Such an understanding can be more easily achieved with the 2D Cylinder Model, which allows for the effect of different geometric parameters to be examined in isolation (Ogawa, Menon, et al., 1993). Furthermore, the computational complexity of the mVAN Model is also the highest, as it takes a considerable amount of computational memory and processing time to simulate the entire macrovasculature, as shown in the next section. The 3D Cylinder Model provides a trade-off between geometrical accuracy and computational requirements, and was investigated as a more practical alternative to the mVAN Model, to potentially make such modelling methods more practical.

#### 4.2.2 Model computational requirements

We also note that high accuracy comes at a cost. The mVAN Model can consume as much as 160 GB of memory and 75 h of computation time for arterial simulations alone). The 3D Cylinder Model, though much simpler (2 GB memory and 108 h computation time), remains much more computationally demanding than 2D Cylinder Model (1 GB memory and 7.5 h computation time) (**Table 2**). According to the discussion above, the 2D Cylinder Model could perform very similarly to the mVAN Model in certain scenarios, and in fact, most S_v,model3_:S_v,model1_ values are lower than 3 (**zoomed window of Fig. S1b**). It is the same for the prediction of correlation coefficients. Thus, in order to select the appropriate simulation model, it is important to consider the error tolerance and the availability of computational resources. Our study is not focused on numerical algorithms, and the computational resources listed here are only meant to provide a comparison and may not be based on the most efficient algorithms or reflect all hardware architectures.

**Table 2.**
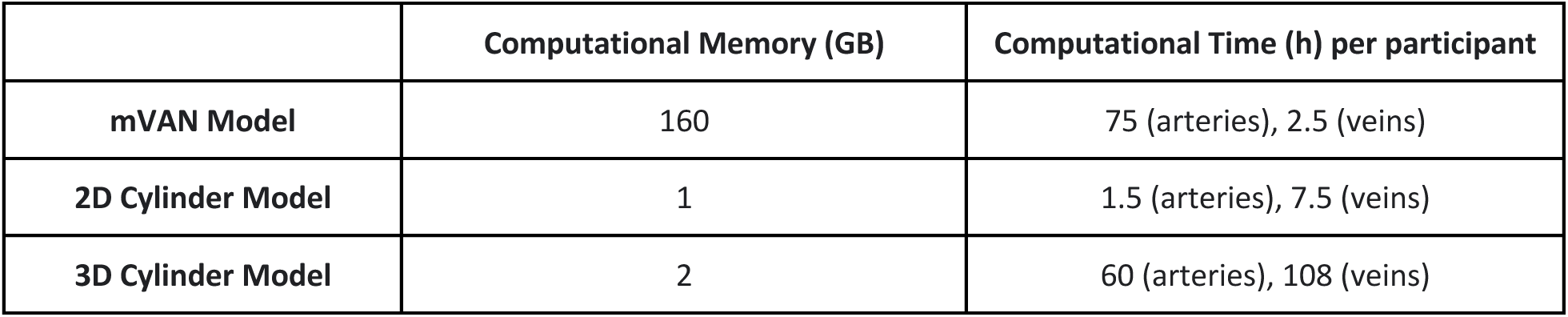
Computational resource for each model.

|  | Computational Memory (GB) | Computational Time (h) per participant |
| --- | --- | --- |
| mVAN Model | 160 | 75 (arteries), 2.5 (veins) |
| 2D Cylinder Model | 1 | 1.5 (arteries), 7.5 (veins) |
| 3D Cylinder Model | 2 | 60 (arteries), 108 (veins) |

#### 4.2.3 Model experimental comparisons

To establish a point of reference for the comparison across the three models, the simulated baseline BOLD signal intensities were compared (**Fig. S1**). The RSFA shifts the comparison from static to dynamic. A general conclusion can be drawn that the mVAN Model provides better predictions than the 2D Cylinder and 3D Cylinder Models for RSFA (**Fig. 6 and 10**). Interestingly, The mVAN Model was able to predict venous-driven RSFA reasonably well. As discussed in the theory section, the mVAN Model has the most realistic geometric representation of macrovasculature, and while the 3D Cylinder Model is less realistic, it still embodies the interactions between vascular orientation and fBV, whereas 2D Cylinder Model does not. Nevertheless, none of the three models is capable of providing satisfactory prediction for arterial RSFA.

Generally, the mVAN Model resulted in higher R^2^ values than the 2D and 3D Cylinder Model for all three forms of macrovascular FC studied (arterial-arterial, arterial-venous, and venous-venous). In addition, the 3D Cylinder Model generally performed better than the 2D Cylinder Model in all cases (**Fig. 6**). Nevertheless, we noted in cases with poor prediction that the simulated values appear to be subdivided into two groups, the first spanning correlation coefficients greater than 0.04, and the second spanning correlation coefficients between 0 and 0.04. The existence of two clusters of correlation coefficients may result from signal intensity outliers as discussed above (see *Model Theoretical Comparison*). In light of the results in **Fig. S1**, the 2D Cylinder Model is likely to produce spurious BOLD signal fluctuation amplitudes with specific vascular orientation and fBV values that may distort the predicted time courses. This may translate into extreme RSFA values and lead to widely fluctuating FC values. On the other hand, both the mVAN Model and the 3D Cylinder Model generated satisfactory predictions for venous-venous correlation, which suggests that the 3D Cylinder Model may be used as a substitute for the mVAN Model.

### 4.3 Theoretical understanding of macrovascular BOLD

According to the classic biophysical model proposed by Ogawa et al. (Ogawa, Menon, et al., 1993), the BOLD signal is dependent on the vascular orientation, size, and oxygenation. θ and fBV are two parameters widely investigated in relation to BOLD signal dependencies (Ogawa, Menon, et al., 1993; Viessmann et al., 2019; Zhong & Chen, 2022). The ΔBOLD metric, calculated based on difference in the BOLD signal corresponding to the two simplified cases of the input signals (variations in fBV for arteries and in Y for veins), was used to demonstrate the dependency of each parameter on the simulated BOLD fluctuations, whose magnitude should be proportional to the RSFA and whose polarity will directly affect the correlation with other BOLD signals from other locations.

In contrast with previous experimental observations at 7 T based primarily on mesoscopic pial veins and the associated extravascular fields (Viessmann et al., 2019), the magnitude of macrovascular ΔBOLD in this work does not consistently decrease with increasing θ, but is rather minimised at θ ∼ 3π/4 (**Fig. 5m,n**), which, according to the original biophysical model (Ogawa, Menon, et al., 1993), leads to zero intravascular magnetic-field offset. The macrovasculature, in contrast to microvasculature, has a higher fraction of intravascular volume, such that its IV contribution cannot be neglected. It is intriguing to note that, although venous ΔBOLD remains positive for all fBV, arterial ΔBOLD changes polarity from negative to positive with an fBV of ∼0.4 (**Fig. 5o,p**). It is possible that a decrease in the extravascular compartment associated with an fBV increase could lead to a dominance of intravascular contribution to the measured BOLD fluctuations. (In the case of arteries, an increase in the intravascular compartment would be expected to increase phase homogeneity, since all intravascular spins in a given voxel would be in phase.) As a result of the change in polarity of the simulated signal, arterial-arterial correlations could potentially be negative with specific sets of fBV, just as the arterial-venous correlation could be positive. These complicated dependencies suggest that multiple factors must be considered simultaneously when modelling macrovascular contribution to both BOLD fluctuation magnitude and correlation, and that simply considering diameter would not suffice for voxel-based modelling (Huck et al., 2023).

However, these are not the only parameters that could affect the simulated BOLD signals. Our previous study demonstrated that even unexpected parameters such as the position of a vessel within a voxel can have an impact on macrovascular BOLD (Sedlacik et al., 2007; Zhong & Chen, 2023) and this can be extended to the impact of partial-volume effects. It is, however, difficult to measure the exact position of vessels within the fMRI voxels in our in-vivo experimental data, so the simulation was not able to include this effect. It is also possible that our results may be affected by other parameters (such as TE, spatial resolution, and the main magnetic field), which are not addressed in this initial study but could be considered in the future.

### 4.4 Experimentally measured macrovascular BOLD: agreement with prediction

#### 4.4.1 Model prediction of macrovascular BOLD standard deviations

The mVAN Model was able to predict venous RSFA reasonably well. However, this is not the case with the other two models. In the 2D Cylinder and the 3D CylinderModels, we observed a cluster of low simulated values that do not fit well into a linear relationship, which suggests unreliable prediction for some of the voxels. The reason could be explained by the same logic presented above in *Model Comparison*, which is that the unrealistic geometry of the 2D Cylinder Model makes its predicted signal fluctuation amplitude deviate (up to 60%) from that of the mVAN Model. Moreover, the RSFA prediction accuracy is highly participant dependent as seen in **Fig. 10**, which may in turn be dependent on the parameters of the acquisition and on the distinct anatomical structures of each participant (e.g., participant positioning and geometry of the macrovasculature). Moreover, different participants may also be affected by varying levels of physiological noise (e.g. heart rate variability, respiratory variability), which further increases the level of inter-subject variability. As well, the effect of motion artefacts cannot be ignored. Different participants may have different levels of motion (either gross head motion or local physiological pulsation) which result in different levels of noise for both rs-fMRI and TOF imaging.

In general, unlike for venous RSFA, none of our models provided satisfactory predictions of arterial RSFA in any of our subjects (**Fig. 10**). Since the diameter of the arteries are constantly changing, it is almost impossible to establish an accurate baseline diameter for use in our simulations. Additionally, the duration of the TOF angiogram acquisition is significantly longer than the arterial variation (the MRA acquisition in the MSC dataset lasted approximately five minutes) (Gordon et al., 2017; Reimers et al., 2018), resulting in a further blurring of the arterial vasculature boundaries. A further challenge is emulating arterial movement in simulation, particularly for the mVAN Model, which requires extremely high resolution and is therefore not practicable. While some state-of-the-art acquisition techniques allow dynamic imaging of arteries, they still only cover one blood vessel rather than the entire vasculature (Varadarajan et al., 2023). Further research to improve the temporal resolution of angiograms will be necessary.

#### 4.4.2 Model prediction of macrovascular BOLD correlation coefficients

The discrepancy between the predicted and the experimental correlation coefficient could be, in part, due to the simulated signal having a much higher signal-to-noise ratio (SNR) than the in-vivo signal. To account for this, we weight the simulated connectivity with the simulated RSFA in order to emulate the in-vivo experimental signals better (Golestani & Goodyear, 2011), as described in the methods section. This approach prevented the FC maps from being dominated by large vessels.

Venous-venous correlation was remarkably well predicted by the mVAN Model, followed by arterial-venous correlation. Similar to the case of RSFA, we were not able to predict arterial-arterial correlation with any of the models. Interestingly, predicted arterial-venous correlations were inverted relative to the experimental values (**Fig. 7d, f**). Arterial BOLD contributions are generally weaker than venous contributions, whereas the angiograms are more sensitive to the fast-flowing arterial blood than the slower moving blood of the neighbouring veins (such as the cavernous sinus, which is in close proximity to the Circle of Willis). Thus, when a single voxel contains both an artery and a vein, one dominating the segmentation and the other dominating the susceptibility effect, the model prediction may be particularly challenging. Certainly, given the poor accuracy of arterial predictions seen in this study, it is clearly more feasible to pivot towards estimating large-vein contributions using our approach. Our R^2^ values are lower than those of the binned predictions reported by Huck et al. (R^2^>0.98) (Huck et al., 2023), but much higher than those of their voxel-wise predictions (R^2^<0.05). The results of our study, although still based on binning, are based only on predicted correlation coefficients from first principles, which should allow better representation of the voxel-wise macrovascular effects. Since Huck et al. also examined other connectivity metrics (low-frequency fluctuation amplitude, regional connectivity inhomogeneity, Hurst exponent, and eigenvector centrality), a direct comparison of these two studies is difficult.

Interestingly, our prediction of correlation coefficients appears to be more accurate than our prediction of RSFA, especially when comparing venous RSFA and venous-venous correlation. There may be several reasons for this. First, measured RSFA likely includes factors that are not incorporated in our model such as multiplicative scaling from receive coil sensitivities, which may scale amplitudes of the fluctuations not but not their timing. Correlation coefficients are normalized quantities and thus are less influenced by these regional scalings. Second, following similar logic, the RSFA and correlation coefficient are also differentially sensitive to partial-volume effects. Third, factors such as vascular position in a voxel also affect signal intensity fluctuations but not necessarily the correlation (Zhong & Chen, 2023). The basic concept stems from the fact that a given voxel may not be able to sample the entire dipole, but the details are beyond the scope of this paper. The same argument can be extended to the relative immunity of correlation coefficients to errors in estimating fBV and orientation from the segmented macrovasculature because of how they are normalized. Such errors will also have a greater impact on the fluctuation magnitude than the temporal features of the fluctuations.

#### 4.4.3 Model prediction of perivascular BOLD effects

In both arteries and veins, our model was unable to predict the in-vivo experimental RSFA at locations farther than one voxel away (∼4 mm) from the macrovasculature. This may correspond to different numbers of voxels depending on the voxel size. A potential explanation for the unsatisfactory predictions could be found in the RSFA maps (**Fig. S6**), which suggest that the macrovascular susceptibility effect may only extend, or be predictable, over a limited distance from the macrovasculature. Consequently, in ROIs with limited macrovascular susceptibility effects, the in-vivo signal may be dominated by other sources of fluctuations (for example, neural activity and physiological noise), which cannot be reflected by the predicted RSFA.

From the perspective of correlations, we found that in-vivo venous-venous correlations, while extremely well predicted by the mVAN Model, also became substantially less well predicted at one voxel away from the macrovasculature. This finding is similar to that of a previous study that found that the contribution of the veins to connectivity decreased with distance from the venous vasculature (Huck et al., 2023). Arterial-arterial correlations were not well predicted by the simulations even in the voxels containing the vessels.

One of the important questions raised by our previous studies is whether perivascular FC is related to EV magnetic susceptibility or to contributions of undetected vessels. As discussed in the previous section, the magnetic susceptibility difference between blood and tissue would generate an extravenous dipolar magnetic field offset around macrovasculature that could extend well beyond the voxel containing the vessel (Ogawa, Menon, et al., 1993). We observed that the susceptibility effect became attenuated rapidly beyond macrovascular voxel (**Fig. S6**), and was unlikely to persist more than 4 mm away from the macrovasculature (even when thresholded a 1% of the maximum RSFA). Our regression results, for both RSFA and correlation coefficients, support this claim. Such would be the case whether the macrovascular R_2_’ perturbations are due to neuronal activity or other physiological processes. Of course, some physiological noise effects are global (Birn et al., 2006; Chang et al., 2009; Tong et al., 2015), however they would not cause the model performance to deteriorate with increasing distance from the macrovasculature. Since the latter is what we see, we propose that EV susceptibility effects are not the sole contributor to perivascular BOLD. There are numerous medium-to-small blood vessels in the extra-macrovascular space that are not detected by the TOF acquisition due to their smaller size and lower flow velocity. Their unmodeled BOLD contributions, which are likely also modulating the BOLD signal behaviour more distal to the microvasculature, may be the main reason that our modeling accuracy deteriorates with increasing distance from the macrovasculature. Our future work will target the improvement of macrovascular detection sensitivity, perhaps through the use of contrast agents (Bernier et al., 2020).

### 4.5 Recommendations

This study aims to demonstrate that modelling the macrovascular BOLD FC (in terms of correlation coefficients) using a biophysical model is feasible, and can provide a post-acquisition means to reduce the macrovascular bias during data analysis and interpretation. This study demonstrates, in concept, that TOF data acquired within rs-fMRI sessions can be used to predict the macrovascular bias, as we suggested in our previous study (Zhong et al., 2023). The use of a full macrovascular VAN (the mVAN Model in this work) demonstrated the highest performance in predicting venous-driven macrovascular FC, and thus should be the starting point for future efforts to eliminate macrovascular contribution on FC. To address the high computational cost of such an approach, pre-computed look-up tables could be developed that encapsulate all possible simulated signals, thereby increasing the accessibility of our approach. However, the prediction of arterial macrovascular BOLD is poor, potentially calling for better methods of capturing arterial dynamics. Lastly, perivascular BOLD contributions by large veins could be modelled reasonably accurately at ∼4 mm beyond the vascular voxel, but not more. This could speak to the need to use more sensitive TOF imaging and higher spatial resolution to model perivascular effects from smaller vessels.

Based on the current results, adjusting correlation coefficients based on simulated correlation coefficients would be the most promising way to reduce the macrovascular effect on resting-state fMRI. That is, the macrovascular cross-correlation coefficients could be accurately predicted by our mVAN Model. It is, however, important to realize that such an FC correction is currently only realizable on the cross-correlation coefficients (instead of Pearson’s correlation, for example) between voxels in the vicinity of large veins. This is because our models do not account for lags between times series and their effects on correlation coefficients. Besides correlations, there may also be other, lag-insensitive approaches for calculating FC, such as regression and independent component analysis (ICA), but adapting our investigation to these other metrics will be the subject of our future work.

### 4.6 Study limitations

This study has several limitations. This report will not discuss the limitations of in-vivo experiments, which are discussed in our previous work (Zhong et al., 2023). We will instead focus on the limitations most relevant to the simulations. The two biggest limitations of the simulations would be simulation completeness limited to TOF sensitivity and simulation feasibility limited by computational demand. Due to the limited sensitivity of the TOF data, we are unable to model the entire macrovasculature, which makes the input macrovasculature imperfect for fully accurate simulations and will reduce prediction performance. Moreover, the accuracy of vascular fBV estimates can be biassed by differences in flow rates across vessels. Additionally, the simulation is based on simple sinusoidal signals as inputs to mimic resting-state fluctuations, which is likely not realistic (for either fBV or Y). Also, some parameters, particularly for the 2D and 3D Cylinder Model, were shown to be important, but were difficult to extract from TOF images, such as the position of the vessel within the fMRI voxel (Zhong & Chen, 2023). Furthermore, there have been various definitions of “macrovasculature” in the literature. Recognizing that not all venous BOLD effects are artifactual, we will adhere to the defintion of “macrovasculature” outlined in (Menon, 2012) In our specific case, “macrovascular” refers to vessels detectable from our TOF MRA data, which are far larger in diameter than the intracortial veins.

The availability of computational resources also limits the feasibility of the simulation. In spite of having more than 180 GB of memory, our server can only support 0.2-mm resolution for the mVAN Model simulation, which is not sufficient for some narrow segments of the macrovasculature. The long computational times for the simulations is another factor, which only allowed us to include four participants within a reasonable timeframe. There is a possibility that this problem may be alleviated by the addition of computational resources in the near future.

Furthermore, the applicability of these results likely depends on scan parameters (including spatial resolution, main magnetic field and temporal resolution) of the MSC dataset, therefore comparisons with previous studies, especially those with higher main magnetic fields, should be made with caution. In practice, predictions should be tailored to different scanning protocols.

## 5 Conclusions

To conclude, we found that the macrovascular FC could be predicted using biophysical modelling based on vascular anatomical information, and that the venous contributions could be more accurately predicted than the arterial contributions. In addition, we found that the models were less reliable in predicting macrovascular RSFA than FC. Moreover, we found that modelling perivascular BOLD effects using simulations is feasible at a distance close to macrovasculature, also more accurately for veins than for arteries. This study paves a path for model-based correction of the macrovascular bias in resting-state and other types of fMRI, the validation of which will be part of our future work.

## Acknowledgments

The authors would like to acknowledge financial support from Canadian Institutes of Health Research and the Canada Research Chairs Program (JJC), funding support from NIH NIBIB grant P41-EB030006, NIMH grant R01-MH111419 and NINDS grant U19-NS123717 (JRP), and funding support from Ydessa Hendeles Graduate Scholarship (XZZ).

## Ethics

The study was approved by Baycrest REB (#11-47).

## Author contributions

Xiaole (Zachary) Zhong: Conceptualization, methodology, software, investigation, writing-original draft preparation. Jonathan R. Polimeni: writing-reviewing and editing. J. Jean Chen: Conceptualization, methodology, software, investigation, writing-original draft preparation, supervision, writing-reviewing and editing.

## Data and Code Availability

All data described in this manuscript was obtained from the Midnight Scan Club (MSC) dataset. The data is available publicly at OpenNeuro. The code will be made available upon request.

## Reference

1. Andrews-Hanna, J. R., Snyder, A. Z., Vincent, J. L., Lustig, C., Head, D., Raichle, M. E., & Buckner, R. L. (2007). Disruption of large-scale brain systems in advanced aging. Neuron, 56(5), 924–935. 10.1016/j.neuron.2007.10.038

2. Aso, T., Sugihara, G., Murai, T., Ubukata, S., Urayama, S.-I., Ueno, T., Fujimoto, G., Thuy, D. H. D., Fukuyama, H., & Ueda, K. (2020). A venous mechanism of ventriculomegaly shared between traumatic brain injury and normal ageing. Brain: A Journal of Neurology, 143(6), 1843–1856. 10.1093/brain/awaa125

3. Bandettini, P. A., Jesmanowicz, A., Wong, E. C., & Hyde, J. S. (1993). Processing strategies for time-course data sets in functional MRI of the human brain. Magnetic Resonance in Medicine: Official Journal of the Society of Magnetic Resonance in Medicine / Society of Magnetic Resonance in Medicine, 30(2), 161–173. 10.1002/mrm.1910300204

4. Bandettini, P. A., & Wong, E. C. (1995). Effects of biophysical and physiologic parameters on brain activation-induced R2* and R2 changes: Simulations using a deterministic diffusion model. International Journal of Imaging Systems and Technology, 6(2-3), 133–152. 10.1002/ima.1850060203

5. Berman, A. J. L., Mazerolle, E. L., MacDonald, M. E., Blockley, N. P., Luh, W.-M., & Pike, G. B. (2018). Gas-free calibrated fMRI with a correction for vessel-size sensitivity. NeuroImage, 169, 176–188. 10.1016/j.neuroimage.2017.12.047

6. Bernier, M., Cunnane, S. C., & Whittingstall, K. (2018). The morphology of the human cerebrovascular system. Human Brain Mapping, 39(12), 4962–4975. 10.1002/hbm.24337

7. Bernier, M., Pfannmoeller, J. P., Bollmann, S., Berman, A. J. L., & Polimeni, A. J. R. (2021, April 30). Modeling the vascular influences on BOLD fMRI using in vivo brain vasculature: incorporating vessel diameter, orientation, and susceptibility. 2021 ISMRM & SMRT Annual Meeting & Exhibition. ISMRM.

8. Bernier, M., Viessmann, O., Ohringer, N., Chen, J. E., Fultz, N. E., Leaf, R. K., Wald, L. L., & Polimeni, J. R. (2020). Human cerebral white-matter vasculature imaged using the blood-Pool contrast agent ferumoxytol: bundle-specific vessels and vascular density. Proc Intl Soc Mag Reson Med, 28, 0016. https://archive.ismrm.org/2020/0016.html

9. Billett, H. H. (1990). Hemoglobin and Hematocrit. In H. K. Walker, W. D. Hall, & J. W. Hurst (Eds.), Clinical Methods: The History, Physical, and Laboratory Examinations. Butterworths. https://www.ncbi.nlm.nih.gov/pubmed/21250102

10. Birn, R. M., Diamond, J. B., Smith, M. A., & Bandettini, P. A. (2006). Separating respiratory-variation-related fluctuations from neuronal-activity-related fluctuations in fMRI. NeuroImage, 31(4), 1536– 1548. 10.1016/j.neuroimage.2006.02.048

11. Biswal, B., Yetkin, F. Z., Haughton, V. M., & Hyde, J. S. (1995). Functional connectivity in the motor cortex of resting human brain using echo-planar MRI. Magnetic Resonance in Medicine, 34(4), 537–541. 10.1002/mrm.1910340409

12. Boxerman, J. L., Hamberg, L. M., Rosen, B. R., & Weisskoff, R. M. (1995). MR contrast due to intravascular magnetic susceptibility perturbations. Magnetic Resonance in Medicine, 34(4), 555–566. 10.1002/mrm.1910340412

13. Chang, C., Cunningham, J. P., & Glover, G. H. (2009). Influence of heart rate on the BOLD signal: the cardiac response function. NeuroImage, 44(3), 857–869. 10.1016/j.neuroimage.2008.09.029

14. Cheng, Y.-C. N., Neelavalli, J., & Haacke, E. M. (2009). Limitations of calculating field distributions and magnetic susceptibilities in MRI using a Fourier based method. Physics in Medicine and Biology, 54(5), 1169–1189. 10.1088/0031-9155/54/5/005

15. Chu, P. P. W., Golestani, A. M., Kwinta, J. B., Khatamian, Y. B., & Chen, J. J. (2018). Characterizing the modulation of resting-state fMRI metrics by baseline physiology. NeuroImage, 173, 72–87. 10.1016/j.neuroimage.2018.02.004

16. Cox, R. W. (1996). AFNI: software for analysis and visualization of functional magnetic resonance neuroimages. Computers and Biomedical Research, an International Journal, 29(3), 162–173. 10.1006/cbmr.1996.0014

17. Fan, A. P., Bilgic, B., Gagnon, L., Witzel, T., Bhat, H., Rosen, B. R., & Adalsteinsson, E. (2014). Quantitative oxygenation venography from MRI phase. Magnetic Resonance in Medicine: Official Journal of the Society of Magnetic Resonance in Medicine / Society of Magnetic Resonance in Medicine, 72(1), 149–159. 10.1002/mrm.24918

18. Fischl, B. (2012). FreeSurfer. NeuroImage, 62(2), 774–781. 10.1016/j.neuroimage.2012.01.021

19. Fracasso, A., Luijten, P. R., Dumoulin, S. O., & Petridou, N. (2018). Laminar imaging of positive and negative BOLD in human visual cortex at 7T. NeuroImage, 164, 100–111. 10.1016/j.neuroimage.2017.02.038

20. Frederick, B., & Tong, Y. (2010, May 1). Physiological noise reduction in BOLD data using simultaneously acquired NIRS data. Proceedings of the 16th Annual Meeting of the Organization for Human Brain Mapping. Annual Meeting of the Organization for Human Brain Mapping, Barcelona.

21. Frederick, B., Tong, Y., Strother, M., Nickerson, L., Lindsey, K., & Donahue, M. (2013). Derivation of Flow Information from a Hypocarbia Challenge Study Using Time Delay Correlation Processing. Proceedings of the 21st Annual Meeting of the International Society for Magnetic Resonance In Medicine, Salt Lake City, UT. https://archive.ismrm.org/2013/0206.html

22. Gagnon, L., Sakadžić, S., Lesage, F., Musacchia, J. J., Lefebvre, J., Fang, Q., Yücel, M. A., Evans, K. C., Mandeville, E. T., Cohen-Adad, J., Polimeni, J. R., Yaseen, M. A., Lo, E. H., Greve, D. N., Buxton, R. B., Dale, A. M., Devor, A., & Boas, D. A. (2015). Quantifying the microvascular origin of BOLD-fMRI from first principles with two-photon microscopy and an oxygen-sensitive nanoprobe. The Journal of Neuroscience: The Official Journal of the Society for Neuroscience, 35(8), 3663–3675. 10.1523/JNEUROSCI.3555-14.2015

23. Gao, J. H., Miller, I., Lai, S., Xiong, J., & Fox, P. T. (1996). Quantitative assessment of blood inflow effects in functional MRI signals. Magnetic Resonance in Medicine, 36(2), 314–319. 10.1002/mrm.1910360219

24. Geerligs, L., Renken, R. J., Saliasi, E., Maurits, N. M., & Lorist, M. M. (2015). A Brain-Wide Study of Age-Related Changes in Functional Connectivity. Cerebral Cortex, 25(7), 1987–1999. 10.1093/cercor/bhu012

25. Goense, J. B. M., & Logothetis, N. K. (2006). Laminar specificity in monkey V1 using high-resolution SE-fMRI. Magnetic Resonance Imaging, 24(4), 381–392. 10.1016/j.mri.2005.12.032

26. Golestani, A.-M., & Goodyear, B. G. (2011). A resting-state connectivity metric independent of temporal signal-to-noise ratio and signal amplitude. Brain Connectivity, 1(2), 159–167. 10.1089/brain.2011.0003

27. Gordon, E. M., Laumann, T. O., Gilmore, A. W., Newbold, D. J., Greene, D. J., Berg, J. J., Ortega, M., Hoyt-Drazen, C., Gratton, C., Sun, H., Hampton, J. M., Coalson, R. S., Nguyen, A. L., McDermott, K. B., Shimony, J. S., Snyder, A. Z., Schlaggar, B. L., Petersen, S. E., Nelson, S. M., & Dosenbach, N. U. F. (2017). Precision Functional Mapping of Individual Human Brains. Neuron, 95(4), 791–807.e7. 10.1016/j.neuron.2017.07.011

28. Huck, J., Jäger, A.-T., Schneider, U., Grahl, S., Fan, A. P., Tardif, C., Villringer, A., Bazin, P.-L., Steele, C. J., & Gauthier, C. J. (2023). Modeling venous bias in resting state functional MRI metrics. Human Brain Mapping. 10.1002/hbm.26431

29. Hundley, W. G., Renaldo, G. J., Levasseur, J. E., & Kontos, H. A. (1988). Vasomotion in cerebral microcirculation of awake rabbits. The American Journal of Physiology, *254*(1 Pt 2), H67–H71. 10.1152/ajpheart.1988.254.1.H67

30. Jenkinson, M., Beckmann, C. F., Behrens, T. E. J., Woolrich, M. W., & Smith, S. M. (2012). FSL. NeuroImage, 62(2), 782–790. 10.1016/j.neuroimage.2011.09.015

31. Jensen, J. H., Helpern, J. A., Ramani, A., Lu, H., & Kaczynski, K. (2005). Diffusional kurtosis imaging: the quantification of non-gaussian water diffusion by means of magnetic resonance imaging. Magnetic Resonance in Medicine, 53(6), 1432–1440. 10.1002/mrm.20508

32. Leeper, B. (2015). Venous Oxygen Saturation Monitoring. In Hemodynamic Monitoring: Evolving Technologies and Clinical Practice. Elsevier Health Sciences. https://play.google.com/store/books/details?id=AbzMBgAAQBAJ

33. Li, Y., Zhang, H., Yu, M., Yu, W., Frederick, B. D., & Tong, Y. (2018). Systemic low-frequency oscillations observed in the periphery of healthy human subjects. Journal of Biomedical Optics, 23(5), 1–11. 10.1117/1.JBO.23.5.057001

34. Mayhew, J. E., Askew, S., Zheng, Y., Porrill, J., Westby, G. W., Redgrave, P., Rector, D. M., & Harper, R. M. (1996). Cerebral vasomotion: a 0.1-Hz oscillation in reflected light imaging of neural activity. NeuroImage, *4*(3 Pt 1), 183–193. 10.1006/nimg.1996.0069

35. Menon, R. S. (2002). Postacquisition suppression of large-vessel BOLD signals in high-resolution fMRI. Magnetic Resonance in Medicine, 47(1), 1–9. 10.1002/mrm.10041

36. Menon, R. S. (2012). The great brain versus vein debate. NeuroImage, 62(2), 970–974. 10.1016/j.neuroimage.2011.09.005

37. Menon, R. S., & Goodyear, B. G. (1999). Submillimeter functional localization in human striate cortex using BOLD contrast at 4 Tesla: implications for the vascular point-spread function. Magnetic Resonance in Medicine: Official Journal of the Society of Magnetic Resonance in Medicine / Society of Magnetic Resonance in Medicine, 41(2), 230–235. 10.1002/(sici)1522-2594(199902)41:2>230::aid-mrm3>3.0.co;2-o

38. Mohamed Yacin, S., Srinivasa Chakravarthy, V., & Manivannan, M. (2011). Reconstruction of gastric slow wave from finger photoplethysmographic signal using radial basis function neural network. Medical & Biological Engineering & Computing, 49(11), 1241–1247. 10.1007/s11517-011-0796-1

39. Ogawa, S., Lee, T. M., & Barrere, B. (1993). The sensitivity of magnetic resonance image signals of a rat brain to changes in the cerebral venous blood oxygenation. Magnetic Resonance in Medicine: Official Journal of the Society of Magnetic Resonance in Medicine / Society of Magnetic Resonance in Medicine, 29(2), 205–210. 10.1002/mrm.1910290208

40. Ogawa, S., Menon, R. S., Tank, D. W., Kim, S. G., Merkle, H., Ellermann, J. M., & Ugurbil, K. (1993). Functional brain mapping by blood oxygenation level-dependent contrast magnetic resonance imaging. A comparison of signal characteristics with a biophysical model. Biophysical Journal, 64(3), 803–812. 10.1016/s0006-3495(93)81441-3

41. Ovadia-Caro, S., Margulies, D. S., & Villringer, A. (2014). The value of resting-state functional magnetic resonance imaging in stroke. Stroke; a Journal of Cerebral Circulation, 45(9), 2818–2824. 10.1161/STROKEAHA.114.003689

42. Pan, W.-J., Billings, J. C. W., Grooms, J. K., Shakil, S., & Keilholz, S. D. (2015). Considerations for resting state functional MRI and functional connectivity studies in rodents. Frontiers in Neuroscience, 9, 269. 10.3389/fnins.2015.00269

43. Ragot, D. M., & Chen, J. J. (2019). Characterizing contrast origins and noise contribution in spin-echo EPI BOLD at 3 T. Magnetic Resonance Imaging, 57, 328–336. 10.1016/j.mri.2018.11.005

44. Rebollo, I., Devauchelle, A.-D., Béranger, B., & Tallon-Baudry, C. (2018). Stomach-brain synchrony reveals a novel, delayed-connectivity resting-state network in humans. eLife, 7. 10.7554/eLife.33321

45. Reimers, A. K., Knapp, G., & Reimers, C.-D. (2018). Effects of Exercise on the Resting Heart Rate: A Systematic Review and Meta-Analysis of Interventional Studies. Journal of Clinical Medicine Research, 7(12). 10.3390/jcm7120503

46. Rivadulla, C., de Labra, C., Grieve, K. L., & Cudeiro, J. (2011). Vasomotion and neurovascular coupling in the visual thalamus in vivo. PloS One, 6(12), e28746. 10.1371/journal.pone.0028746

47. Salomir, R., de Senneville, B. D., & Moonen, C. T. W. (2003). A fast calculation method for magnetic field inhomogeneity due to an arbitrary distribution of bulk susceptibility. Concepts in Magnetic Resonance, 19B(1), 26–34. 10.1002/cmr.b.10083

48. Sassaroli, A., Pierro, M., Bergethon, P. R., & Fantini, S. (2012). Low-frequency spontaneous oscillations of cerebral hemodynamics investigated with near-infrared spectroscopy: A review. IEEE Journal of Selected Topics in Quantum Electronics, 18(4), 1478–1492. 10.1109/jstqe.2012.2183581

49. Sedlacik, J., Rauscher, A., & Reichenbach, J. R. (2007). Obtaining blood oxygenation levels from MR signal behavior in the presence of single venous vessels. Magnetic Resonance in Medicine: Official Journal of the Society of Magnetic Resonance in Medicine / Society of Magnetic Resonance in Medicine, 58(5), 1035–1044. 10.1002/mrm.21283

50. Segebarth, C., Belle, V., Delon, C., Massarelli, R., Decety, J., Le Bas, J. F., Décorps, M., & Benabid, A. L. (1994). Functional MRI of the human brain: predominance of signals from extracerebral veins. Neuroreport, 5(7), 813–816. 10.1097/00001756-199403000-00019

51. Shin, W., Gu, H., & Yang, Y. (2009). Fast high-resolution T1 mapping using inversion-recovery Look-Locker echo-planar imaging at steady state: optimization for accuracy and reliability. Magnetic Resonance in Medicine, 61(4), 899–906. 10.1002/mrm.21836

52. Spees, W. M., Yablonskiy, D. A., Oswood, M. C., & Ackerman, J. J. (2001). Water proton MR properties of human blood at 1.5 Tesla: magnetic susceptibility, T(1), T(2), T*(2), and non-Lorentzian signal behavior. Magnetic Resonance in Medicine: Official Journal of the Society of Magnetic Resonance in Medicine / Society of Magnetic Resonance in Medicine, 45(4), 533–542. 10.1002/mrm.1072

53. Srivastava, P., Fotiadis, P., Parkes, L., & Bassett, D. S. (2022). The expanding horizons of network neuroscience: From description to prediction and control. NeuroImage, 258(119250), 119250. 10.1016/j.neuroimage.2022.119250

54. Stanley, O. W., Kuurstra, A. B., Klassen, L. M., Menon, R. S., & Gati, J. S. (2021). Effects of phase regression on high-resolution functional MRI of the primary visual cortex. NeuroImage, 227, 117631. 10.1016/j.neuroimage.2020.117631

55. Tang, C., Blatter, D. D., & Parker, D. L. (1993). Accuracy of phase-contrast flow measurements in the presence of partial-volume effects. Journal of Magnetic Resonance Imaging, 3(2), 377–385. 10.1002/jmri.1880030213

56. Thayer, J. F., Ahs, F., Fredrikson, M., Sollers, J. J., 3rd, & Wager, T. D. (2012). A meta-analysis of heart rate variability and neuroimaging studies: implications for heart rate variability as a marker of stress and health. Neuroscience and Biobehavioral Reviews, 36(2), 747–756. 10.1016/j.neubiorev.2011.11.009

57. Tong, Y., Bergethon, P. R., & Frederick, B. D. (2011). An improved method for mapping cerebrovascular reserve using concurrent fMRI and near-infrared spectroscopy with Regressor Interpolation at Progressive Time Delays (RIPTiDe). NeuroImage, 56(4), 2047–2057. 10.1016/j.neuroimage.2011.03.071

58. Tong, Y., & Frederick, B. D. (2010). Time lag dependent multimodal processing of concurrent fMRI and near-infrared spectroscopy (NIRS) data suggests a global circulatory origin for low-frequency oscillation signals in human brain. NeuroImage, 53(2), 553–564. 10.1016/j.neuroimage.2010.06.049

59. Tong, Y., & Frederick, B. D. (2012). Concurrent fNIRS and fMRI processing allows independent visualization of the propagation of pressure waves and bulk blood flow in the cerebral vasculature. NeuroImage, 61(4), 1419–1427. 10.1016/j.neuroimage.2012.03.009

60. Tong, Y., & Frederick, B. D. (2014). Studying the Spatial Distribution of Physiological Effects on BOLD Signals Using Ultrafast fMRI. Frontiers in Human Neuroscience, 8, 196. 10.3389/fnhum.2014.00196

61. Tong, Y., Hocke, L. M., Fan, X., Janes, A. C., & Frederick, B. D. (2015). Can apparent resting state connectivity arise from systemic fluctuations? Frontiers in Human Neuroscience, 9, 285. 10.3389/fnhum.2015.00285

62. Tong, Y., Hocke, L. M., & Frederick, B. D. (2011). Isolating the sources of widespread physiological fluctuations in functional near-infrared spectroscopy signals. Journal of Biomedical Optics, 16(10), 106005. 10.1117/1.3638128

63. Tong, Y., Hocke, L. M., & Frederick, B. D. (2014). Short repetition time multiband echo-planar imaging with simultaneous pulse recording allows dynamic imaging of the cardiac pulsation signal. Magnetic Resonance in Medicine: Official Journal of the Society of Magnetic Resonance in Medicine / Society of Magnetic Resonance in Medicine, 72(5), 1268–1276. 10.1002/mrm.25041

64. Tong, Y., Lindsey, K. P., & deB Frederick, B. (2011). Partitioning of physiological noise signals in the brain with concurrent near-infrared spectroscopy and fMRI. Journal of Cerebral Blood Flow and Metabolism: Official Journal of the International Society of Cerebral Blood Flow and Metabolism, 31(12), 2352–2362. 10.1038/jcbfm.2011.100

65. UIudag, K., Müller-Bierl, B., & Ugurbil, K. (2009). An integrative model for neuronal activity-induced signal changes for gradient and spin echo functional imaging. NeuroImage, 47, S56. 10.1016/s1053-8119(09)70204-6

66. Uludağ, K., & Blinder, P. (2018). Linking brain vascular physiology to hemodynamic response in ultra-high field MRI. NeuroImage, 168, 279–295. 10.1016/j.neuroimage.2017.02.063

67. van den Heuvel, M. P., & Hulshoff Pol, H. E. (2010). Exploring the brain network: a review on resting-state fMRI functional connectivity. European Neuropsychopharmacology: The Journal of the European College of Neuropsychopharmacology, 20(8), 519–534. 10.1016/j.euroneuro.2010.03.008

68. Varadarajan, D., Wighton, P., Chen, J., Proulx, S., Frost, R., van der Kouwe, A., Berman, A., & Polimeni, J. R. (2023, May 9). Measuring individual vein and artery BOLD responses to visual stimuli in humans with multi-echo single-vessel functional MRI at 7T. 2023 ISMRM & ISMRT Annual Meeting & Exhibition. ISMRM, Toronto.

69. Vazquez, T., Sañudo, J. R., Carretero, J., Parkin, I., & Rodríguez-Niedenführ, M. (2013). Variations of the radial recurrent artery of clinical interest. Surgical and Radiologic Anatomy: SRA, 35(8), 689–694. 10.1007/s00276-013-1094-4

70. Vemuri, P., Jones, D. T., & Jack, C. R., Jr. (2012). Resting state functional MRI in Alzheimer’s Disease. Alzheimer’s Research & Therapy, 4(1), 2. 10.1186/alzrt100

71. Viessmann, O., Scheffler, K., Bianciardi, M., Wald, L. L., & Polimeni, J. R. (2019). Dependence of resting-state fMRI fluctuation amplitudes on cerebral cortical orientation relative to the direction of B0 and anatomical axes. NeuroImage, 196, 337–350. 10.1016/j.neuroimage.2019.04.036

72. Wise, R. G., Ide, K., Poulin, M. J., & Tracey, I. (2004). Resting fluctuations in arterial carbon dioxide induce significant low frequency variations in BOLD signal. NeuroImage, 21(4), 1652–1664. 10.1016/j.neuroimage.2003.11.025

73. Yu, X., Glen, D., Wang, S., Dodd, S., Hirano, Y., Saad, Z., Reynolds, R., Silva, A. C., & Koretsky, A. P. (2012). Direct imaging of macrovascular and microvascular contributions to BOLD fMRI in layers IV–V of the rat whisker–barrel cortex. NeuroImage, 59(2), 1451–1460. 10.1016/j.neuroimage.2011.08.001

74. Zhang, X., Petersen, E. T., Ghariq, E., De Vis, J. B., Webb, A. G., Teeuwisse, W. M., Hendrikse, J., & van Osch, M. J. P. (2013). In vivo blood T(1) measurements at 1.5 T, 3 T, and 7 T. Magnetic Resonance in Medicine, 70(4), 1082–1086. 10.1002/mrm.24550

75. Zhong, X., & Chen, J. J. (2022, April 22). The dependence of the resting-state macrovascular fMRI signal power on vascular volume and orientation: A simulation study. 2022 Joint Annual Meeting ISMRM-ESMRMB & ISMRT 31st Annual Meeting. ISMRM, London.

76. Zhong, X., & Chen, J. J. (2023, May 19). The dependence of the macrovascular transverse R2’ relaxation and resultant BOLD fMRI signal on vascular position: A simulation study. 2023 ISMRM & ISMRT Annual Meeting & Exhibition. ISMRM, Toronto.

77. Zhong, X., Tong, Y., & Chen, J. J. (2023). Assessment of the macrovascular contribution to resting-state fMRI functional connectivity at 3 Tesla. In bioRxiv (p. 2023.10.19.563131). 10.1101/2023.10.19.563131

