## Supplemental Figures for "Predicting the macrovascular contribution to resting-state fMRI functional connectivity at 3 Tesla: A model-informed approach"

### Signal intensities predicted using the three models

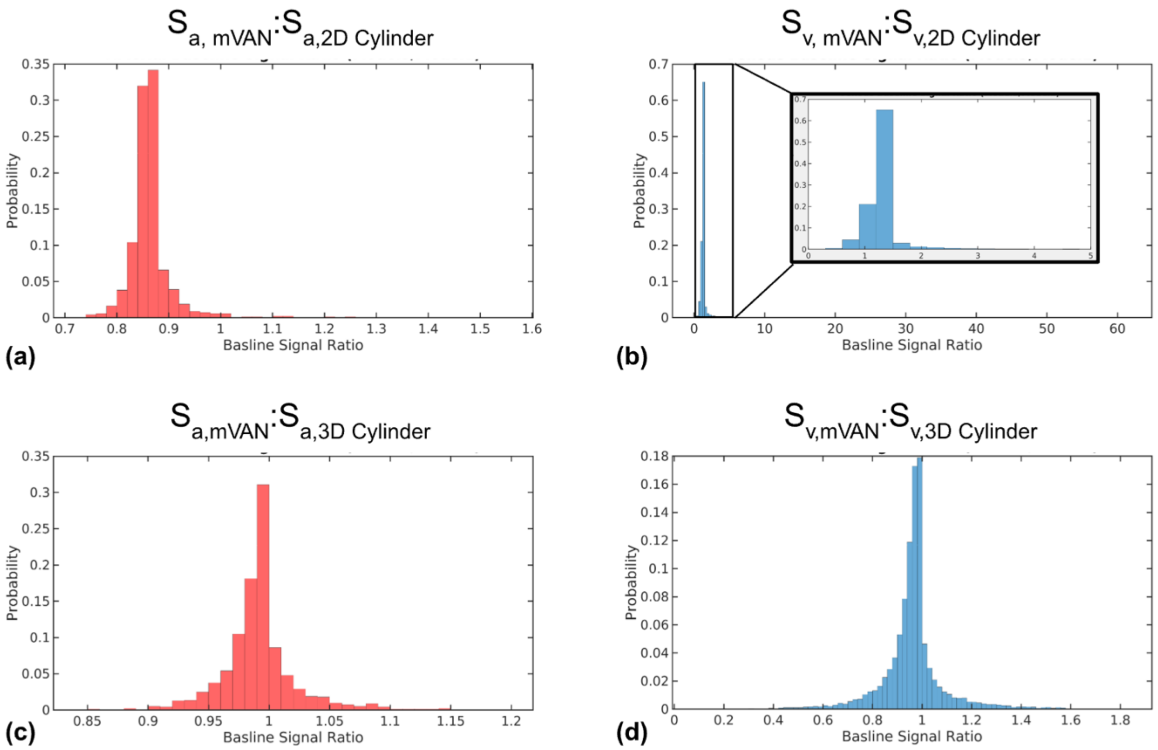


**Figure S1. Signal magnitude comparisons across the three simulation models.** $S_{a,mVAN}$:$S_{a,2D\ Cylinder}$ (a), $S_{v,mVAN}$:$S_{v,2D\ Cylinder}$ (b), $S_{a,mVAN}$:$S_{a,3D\ Cylinder}$ (c) $S_{vmVAN}$:$S_{v,3D\ Cylinder}$ (d). In (b), the histogram spans values up to 60 (unlike in the other panels), but the majority of data points are within a range from 0 to 5.

To compare the three models, we first compared the predicted BOLD signal intensities obtained from them (**Fig. S1**), as this provides a basic indication of how 2D and 3D Cylinder Models differ from the more realistic mVAN model. The comparison was conducted using ratios of pairwise predicted BOLD signal intensities (between all models and mVAN Model), with a ratio of 1 being interpreted as equivalence to mVAN Model. These are based on baseline BOLD intensities (without BOLD signal fluctuations) generated using the default parameters listed in **Table 1**. As shown in **Fig. S1**, the predicted arterial-signal magnitude ratios between mVAN Model and 2D Cylinder Model ($S_{a,mVAN}$:$S_{a,2D\ Cylinder}$) ranged between 0.7 and 1.3 (**Fig. S1a**), whereas venous signal magnitude ratios ($S_{v,mVAN}$:$S_{v,2D\ Cylinder}$) extended to as high as 60 (**Fig. S1b**). While the majority of ratio values are between 0.5 and 3 (as shown in the zoomed inset), the range of ratios between mVAN Model and 2D Cylinder Model far exceeds that of the ratio between mVAN Model and 3D Cylinder Model. These latter ratios ranged between 0.85 and 1.15 for arterial signals ($S_{a,mVAN}$:$S_{a,3D\ Cylinder}$) (**Fig. S1c**) and 0.4 and 1.6 for venous signals ($S_{v,mVAN}$:$S_{v,3D\ Cylinder}$) (**Fig. S1d**).

For veins of the same size and orientation, mVAN Model produced higher BOLD intensities than 3D Cylinder Model ($S_{v,mVAN}$:$S_{v,3D\ Cylinder}$) ranged between 0.4 and 1.6) (**Fig. S1d**), while the intensities were more similar for arteries ($S_{a,mVAN}$:$S_{a,3D\ Cylinder}$) ranged between 0.85 and 1.15) (**Fig. S1c**). The trend continued onto the comparison with the mVAN Model, where the 2D Cylinder Model's venous BOLD intensities were as high as 60 times those of the mVAN Model (**Fig. S1b**). Thus, the 2D and 3D Cylinder Models are not from being equivalent. It should also be noted that most $S_{v,mVAN}$:$S_{v,4D\ Cylinder}$ are below 3 (inset of **Figure S1b**), which suggests that in most cases, venous BOLD predicted using 3D Cylinder Model is reasonably close to those of mVAN Model. Even though these results only reflect a static BOLD effect rather than a dynamic effect, these differences between models may well carry over to dynamic BOLD.

### RSFA predicted using the three models

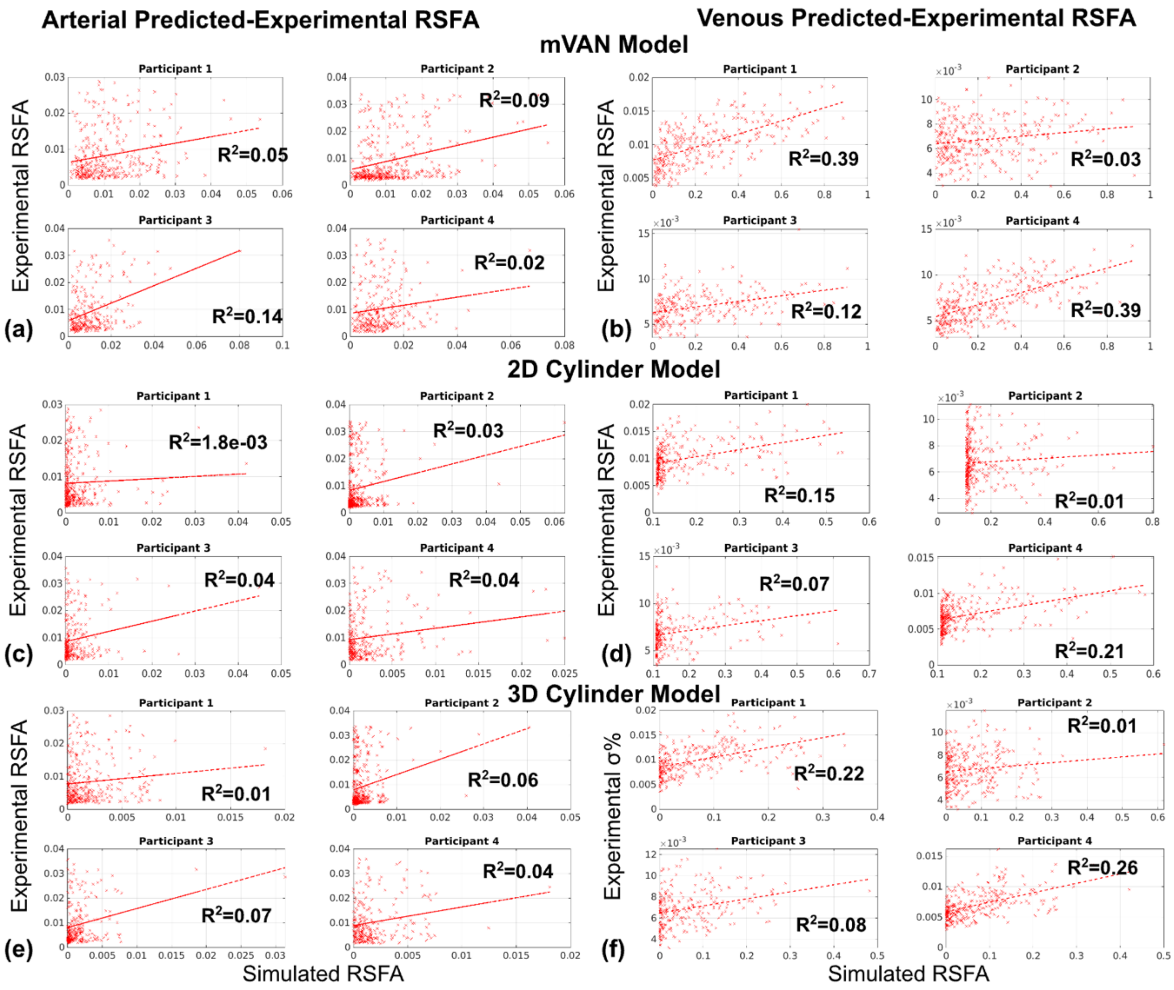


**Figure S2. Data from all subjects: regression between the in-vivo experimental and predicted RSFA for mVAN Model (a,b), 2D Cylinder Model (c,d) and 3D Cylinder Model (e,f).** Arterial and venous results are shown in the left and right halves of the figure, respectively. Each dot represents a bin average.

For mVAN Model, the predicted RSFA exhibited lower predictability for arterial RSFA ($R^2$: 0.02–-0.14) (**Fig. S3a**) and higher for venous RSFA ($R^2$: 0.03–-0.39) (**Fig. S3b**). As with the above models, there is a high degree of inter-subject variability in the simulation-experimental agreement in RSFA. It is still possible to observe large clusters of experimental BOLD RSFA for low values of predicted RSFA, but mainly for arteries and not very strong for veins, indicating some improvement in the venous RSFA prediction.

In all linear fits for 2D Cylinder Model, the RSFA between predicted and in-vivo data were positive, but the goodness of fit was limited. For arteries, $R^2$ ranged between 1.8e−03 to 0.04 (**Fig. S3c**), and for veins, $R^2$ ranged between 0.01 and 0.21 (**Fig. S3d**). For both arteries and veins, the low goodness of fit may be attributed to a large cluster of experimental BOLD RSFA values in voxels that exhibit a low predicted RSFA.

Despite an increase in the maximum $R^2$ to 0.07 fpr 3D Cylinder Model, the accuracy of the prediction of arterial RSFA was still lower than that of veins (**Fig. S3e**), which was associated with positive and higher predictability with $R^2$ between 0.01 to 0.26 (**Fig. S3f**). These data also demonstrate high inter-subject variability in the simulation-experimental agreement in RSFA. Still, for both arteries and veins, the low goodness of fit may be attributed to a large cluster of experimental RSFA corresponding to a low predicted RSFA.

### Correlation coefficients predicted using the three models

#### mVAN Model

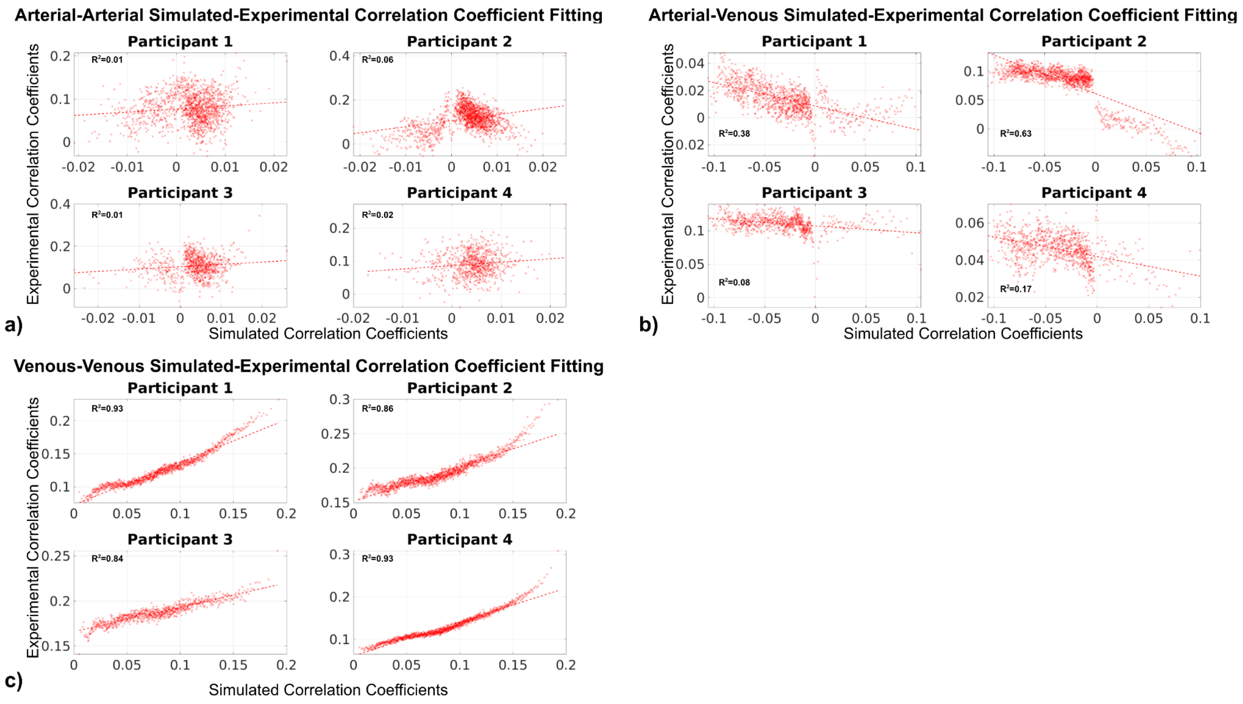


**Figure S3. Regression between the in-vivo experimental and predicted correlation coefficients (mVAN Model).** Illustration of the vascular map model (**Fig. 4c**). Fitting for (a) arterial-arterial correlations, (b) arterial-venous correlations and (c) venous-venous correlations. Each dot represents a bin average.

The predicted correlation coefficients based on mVAN Model show the highest predictiveness for venous-venous correlation coefficients ($R^2$: 0.84–0.93) (**Fig. S3c**), followed by arterial-venous correlation coefficients ($R^2$: 0.08–0.63) (**Fig. S3b**), and with the lowest predictability, arterial-arterial correlation coefficients ($R^2$: 0.01–0.06), (**Fig. S3a**). Additionally, $R^2$ values showed lower inter-subject variability in arterial-arterial and venous-venous correlations.

### 2D Cylinder Model

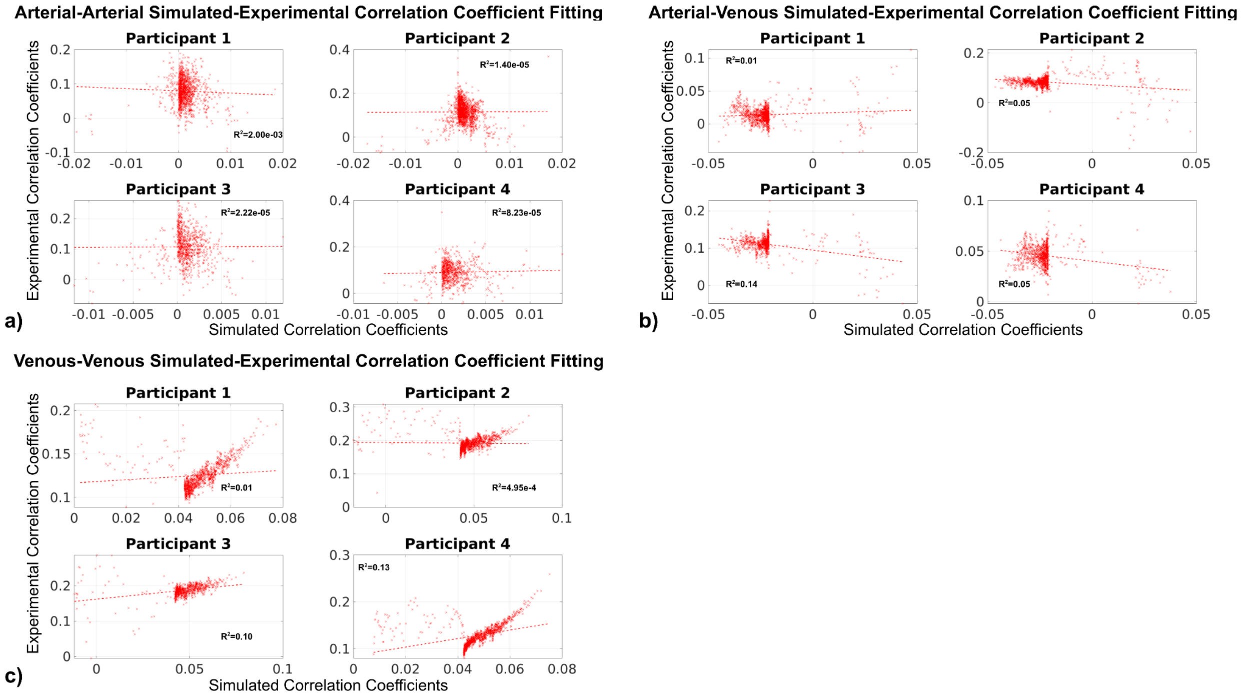


**Figure S4. Regression between the in-vivo experimental and predicted correlation coefficients (2D Cylinder Model).** Illustration of the two-dimensional infinite cylinder model (**Fig. 4a**), with the symbols defined in Table 1. Regression lines for (a) arterial-arterial correlations, (b) arterial-venous correlation, and (c) venous-venous correlation. Each dot represents a bin average.

As expected, for 2D Cylinder Model, most fits showed significant positive correlations between predicted and in-vivo correlation coefficients. However, the goodness of fit was low, with the arterial-arterial correlation $R^2$ being the lowest (< 0.01) (**Fig. S4a**). The arterial-venous correlation $R^2$ was between 0.01 and 0.14 (**Fig. S4b**), and the venous-venous correlation $R^2$ was between 4.9e-3 and 0.13 (**Fig. S4c**). We also noted, especially in **Fig. S4c**, that the data points appear to subdivide into two groups, one spanning correlation coefficients of > 0.04, and the other one spanning the area between 0 and 0.04. As our models do not make provisions for two regimes of correlations, and the ground-truth correlations are unknown in practice, both groups of data were fit to a single linear model.

### 3D Cylinder Model

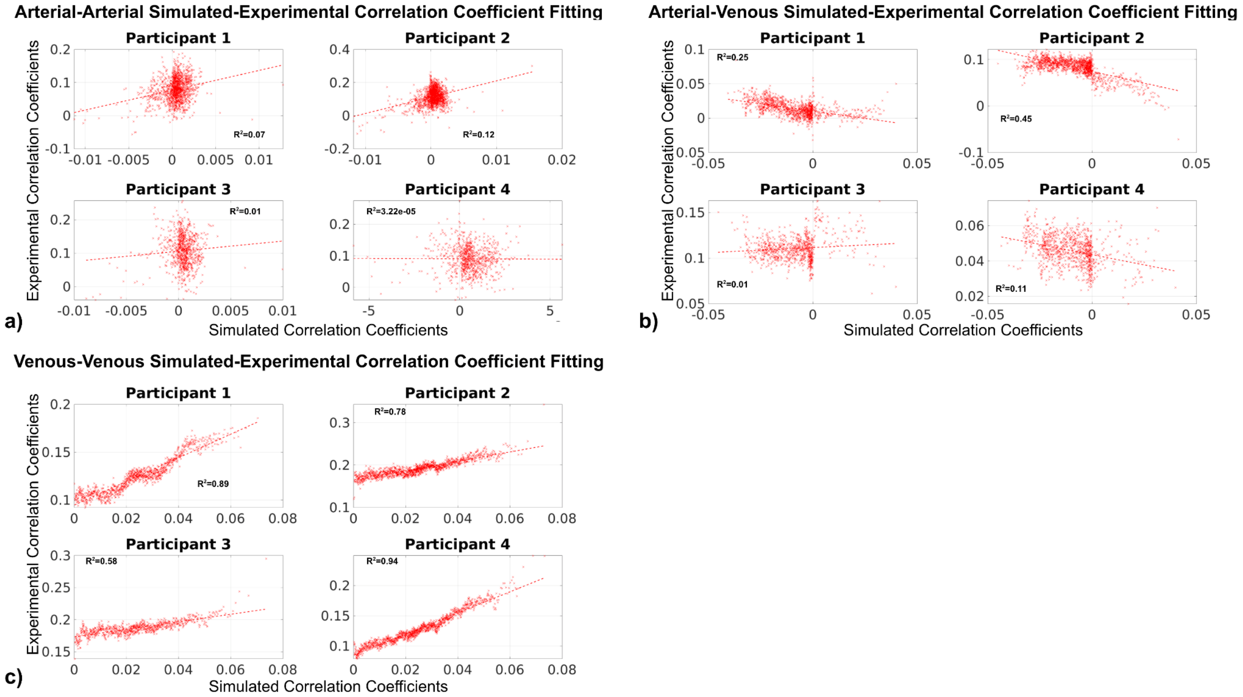


**Figure S5. Regression between the in-vivo experimental and predicted correlation coefficients (3D Cylinder Model).** Illustration of the three-dimensional infinite cylinder model, with the symbols defined in Table 1. Fitting for (a) the arterial-arterial correlation, (b) arterial-venous correlations (c) and venous-venous correlation. Each dot represents a bin average.

For 3D Cylinder Model, despite an increase in the maximum $R^2$ to 0.12, the accuracy of the prediction of arterial-arterial correlations was still relatively low (**Fig. S5a**). The predicted arterial-venous correlations are negatively associated with the experimental ones, with $R^2$ ranging between 9.7e-3 and 0.45 for the four data sets (**Fig. S5b**). However, the predicted venous-venous correlations were significantly and positively associated with experimental ones, with $R^2$ between 0.58 and 0.93 (**Fig. S5c**). These data also demonstrate high inter-subject variability in the simulation-experimental agreement in correlation coefficients.

### Predicted perivascular contributions to GM BOLD signal standard deviations

**Figure S6** summarizes the simulated RSFA maps. To highlight the main trends in the RSFA spatial patterns, RSFA maps from a representative dataset were thresholded at cut-offs of 1% and 10% of the maximum RSFA of the dataset, representing 10 dB and 20 dB below the maximum RSFA. Using the 1% of the maximum RSFA threshold, both arterial and venous simulated signals demonstrated vascular effects compared to extravascular ROIs (**Fig. S6e,f**), but were mainly located one voxel away from the original vascular maps (**Fig. S6a,c**). When the cut-off was increased to 10% of the maximum RSFA, both arterial and venous simulated signals showed minimal extravascular effects (**Fig. S6b,d**).

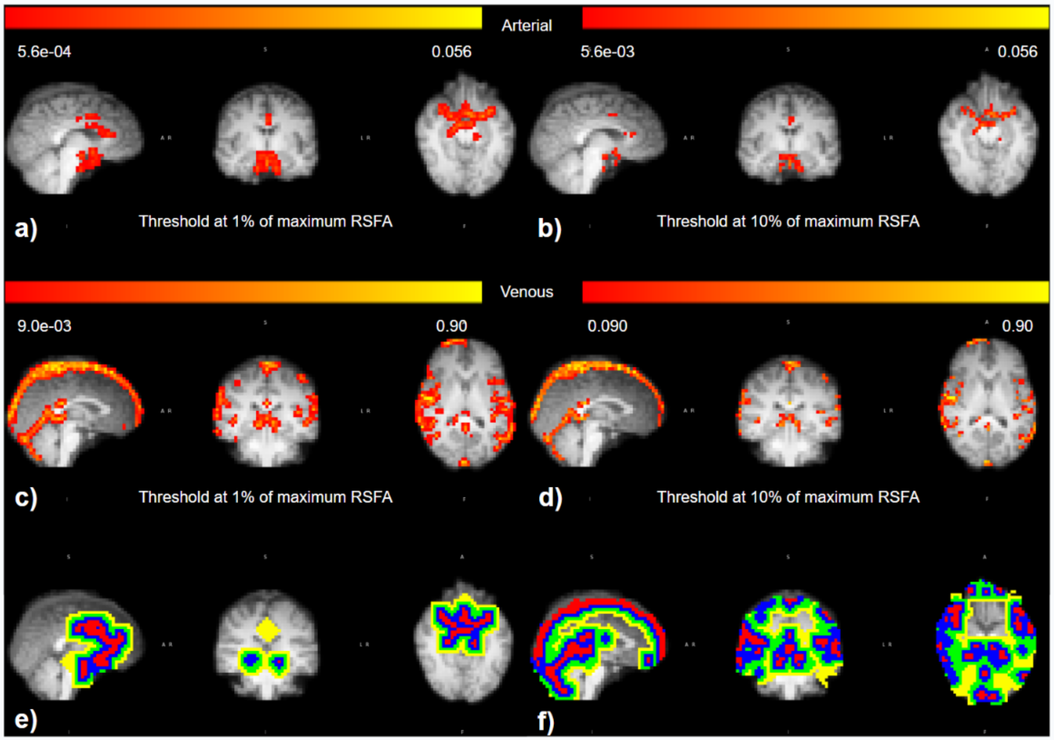


**Figure S6. Maps for normalized RSFA from all experimental data sets (mVAN Model).** An illustration of 1% of the maximum of RSFA cut-off maps (a,b) and 10% of the maximum of RSFA cut-off maps (c,d). Dilated vascular masks are shown for both arterial and venous, with colour coding as: red - voxel containing vasculature, blue: - one voxel away, green: - two voxels away, and yellow: - three voxels away (e,f).

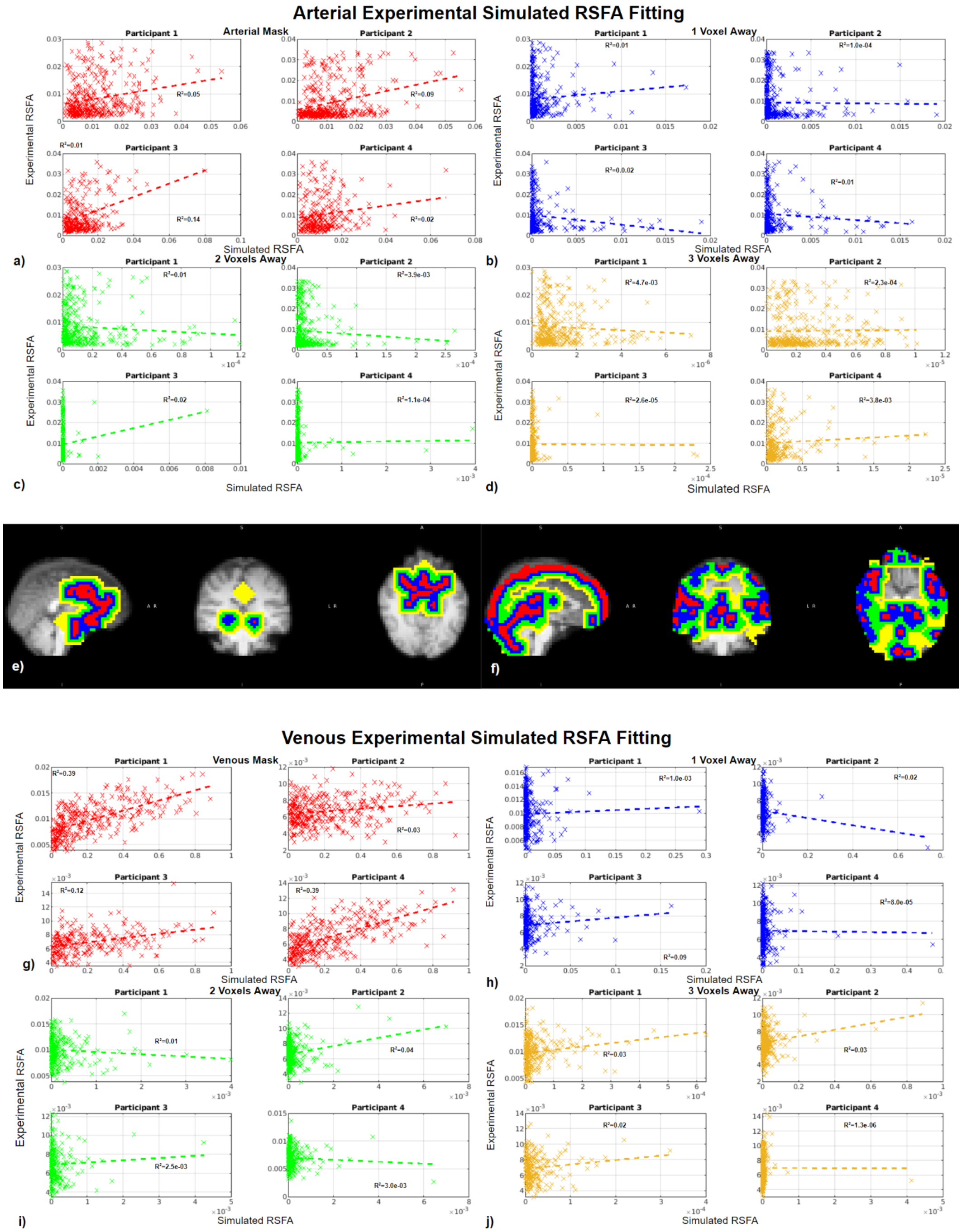


**Figure S7. Dependence of perivascular BOLD standard deviation on distance from vasculature: simulation vs. experimental results (mVAN Model).** An illustration of venous RSFA (a-d) and arterial RSFA (g-j). Perivascular masks are shown for both arterial and venous, with colour coding as: red - containing vessels, blue - one voxel away from the vascular voxel, green - two voxels away, and yellow -three voxels away (e,f).

In the fitting of the perivascular experimental and simulated RSFA for the arterial and venous cases, different patterns are observed with increasing distance from the macrovasculature (**Fig. S7**). Nonetheless, the $R^2$ values still showed relatively low goodness of fit for all arterial macrovascular and perivascular ROIs (all below 0.2) (**Fig. S7a-d**). On the other hand, the $R^2$ values for the venous RSFA could reach 0.39 for certain data sets, and decreased with increasing distance (number of voxels) from the vascular mask. At one voxel away, however, the venous $R^2$ decreased to only 8.0e-05 to 0.02 (**Fig. S7i**) and remained low for two ($R^2$: 2.5e-3 to -0.04) and three voxels ($R^2$: 1.3e-06 to -0.03) away from the macrovascular voxels.

### Predicted perivascular contributions to GM BOLD signal correlations

In the regression of the predicted and measured correlation coefficients in perivascular tissue, different patterns are observed with increasing distance from the macrovasculature in arterial and venous cases (**Fig. S8**). Overall, the $R^2$ values still showed relatively low goodness of fit for all arterial macrovascular and perivascular ROIs (all below 0.2) (**Fig. S8a-d**). However, the $R^2$ values for the venous-venous correlation were as high as 0.93 for certain data sets and decreased with increasing distance (number of voxels) from the vascular mask. The perivascular $R^2$ values for venous-venous correlations, while lower than for the vascular ROI, remained high (ranging from 0.08 to 0.57) at one voxel away from the vascular boundary (**Fig. S8h**). At two voxels away, however, the venous-venous $R^2$ decreased to only 8.3e−05 to 0.12 (**Fig. S8i**). The $R^2$ value (ranging from 7.0e-3 to 0.17) increased again for the three voxels away mask (compared to a distance of two voxels)(**Fig. S8j vs. Fig. S8i**).

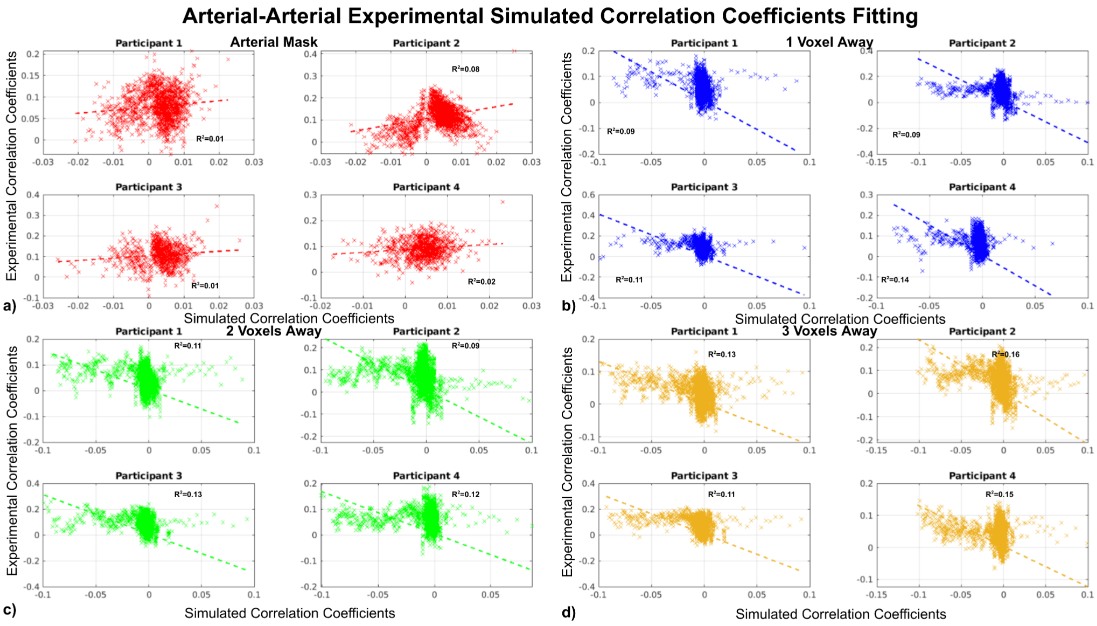

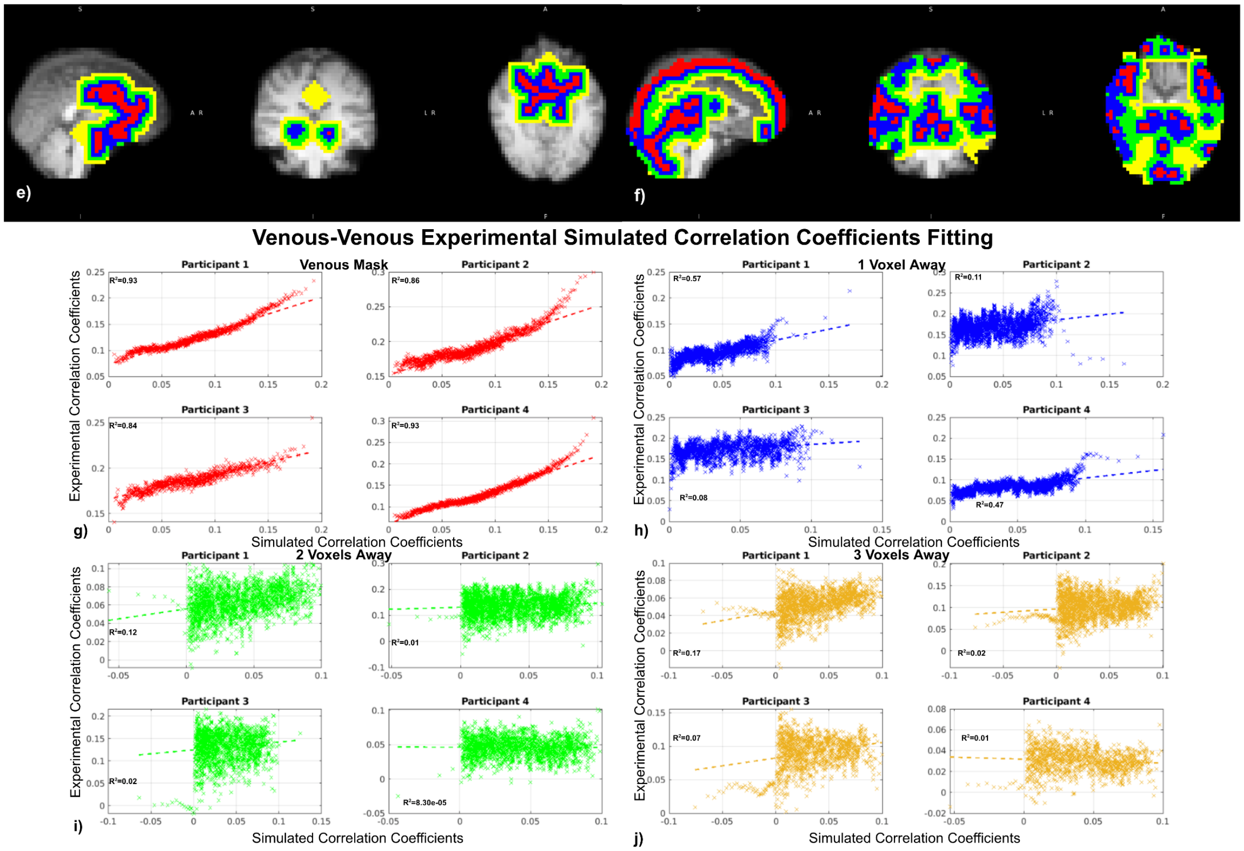


**Figure S8. Dependence of perivascular correlation coefficients on distance from vasculature: simulation vs. experimental results (mVAN Model).** An illustration of venous-venous correlation (a-d) and arterial-arterial correlation (g-j). Perivascular masks are shown for both arterial and venous, with colour coding as: red - intravascular, blue - one voxel away from the vascular voxel, green - two voxels away, and yellow - three voxels away (e,f).
